# Complex compositional and functional diversity of venom metalloproteinases in African puff adders

**DOI:** 10.1101/2024.05.31.596867

**Authors:** Mark C. Wilkinson, Cassandra M. Modahl, Anthony Saviola, Laura-Oana Albulescu, Frank-Leonel Tianyi, Robert A. Harrison, Nicholas R. Casewell

## Abstract

The puff adder (*Bitis arietans*) is a highly venomous viper responsible for many fatalities in Africa, yet there have been few comprehensive analyses of its venom proteins, particularly of the proteases that play a key role in pathology of envenoming. To address this, we have isolated, identified and characterised the bioactivity of the venom metalloproteases of puff adders obtained from a wide range of sources. Prominent in all venoms was an SVMP PI, derived from a PII precursor. This protein existed in either of two forms: non-glycosylated (21 kDa) or glycosylated with either one (26 kDa) or two N-glycans (30 kDa). All the venoms we tested here were found to contain either one or the other form: none had both. The 21 kDa form proved to be highly potent, with alpha-, beta- and in some cases gamma-fibrinogenase activities and were very destructive towards laminin. Prothrombin and Factor X were also extensively degraded by the 21 kDa SVMP, but in neither case did this result in generation of the respective active forms of these clotting factors. In contrast, the two-glycan forms were markedly less active against all of these substrates. The one-glycan form isolated from a Kenyan venom possessed activities that was intermediate between the non- and two-glycan forms. Because of the predominance and ubiquity of these SVMPs in puff adders, and their undoubted clinical significance, we propose to name them the arilysins. The SVMP PIII content of the puff adder venoms was, atypically for African vipers, quite low. In some Kenyan venoms, however, there was an abundant SVMP PIII, with strong gelatinase activity. This protein possesses an unusual oligomeric structure, being a 140 kDa homodimer (c.f. SVMPIII-c) but without the disulphide bonds that normally hold the monomers together in this class of SVMP. This diversity in venom metalloprotease activities is discussed with reference to the potential implications on the pathology of envenomation and the development of therapeutic interventions.

## 1. Introduction

The puff adder (*Bitis arietans*) is found throughout much of the African continent and western Arabia and is considered to be one of the most medically important snakes in this region, accounting for a high proportion of the 200-300,000 annual envenomings in sub-Saharan Africa (Chippaux, 2011). Despite this, there have been too few studies on the clinical effects of puff adder envenoming. Most notable of these is that of Warrell et al. (1975) who recorded details of 10 Nigerian patients envenomed by puff adders and more recently Tianyi et al (2024) carried out a thorough case-series of puff adder bites treated at two primary healthcare facilities in Kenya, complemented with a scoping review of published cases of individual bites. Both studies found local and systemic clinical features of puff adder bites to be diverse, though inconsistent, and included pain, swelling, necrosis, haemorrhage, coagulopathy, thrombocytopoenia, fever, and hypotension/shock.

These diverse pathologies of puff adder envenoming are due to the proteins in the injected venom, whose composition is broadly similar to that of other vipers (Casewell et al., 2014; Currier et al., 2010; Fasoli et al., 2010, Dawson et al. 2024). The three most abundant protein classes in this venom: C-type lectin-like proteins (CLPs), snake venom metalloproteases (SVMPs) and snake venom serine proteases (SVSPs) are highly likely to be those responsible for the bulk of the venom-induced pathologies. Owing to their well-documented effects on coagulation and platelet physiology (Arlinghaus and Eble, 2012; Morita, 2005), the CLPs will be partially responsible for two of the known systemic effects of puff adder envenomating: thrombocytopoenia and coagulopathy. It is likely, however, that the SVMPs and SVSPs will also play a major role in coagulopathy and are undoubtedly responsible for other commonly observed hemotoxic effects such as spontaneous haemorrhage and hypotension. A few early studies identified EDTA-inhibited haemorrhagic components in puff adder venom, but only in two cases has an SVMP with this activity been isolated. Thus, Van der Walt et al., (1972) purified and characterised a 21 kDa SVMP, and Omori-Satoh et al., (1995) found two distinct haemorrhagic SVMPs with molecular weights of 68 and 75 kDa. Hypotension in patients with puff adder envenoming can be an indirect consequence of haemorrhage and also of the direct action of adenosine (Graham et al., 2005) and bradykinin potentiating peptides (Kodama et al., 2015), kallikrein-like (kinin-releasing) SVSPs (Megale et al., 2018; Sekoguchi et al., 1986; Nikai et al., 1993). There is also evidence that SVMPs in puff adder venom may contribute to the hypotensive action through generation of the vasodilator angiotensin 1-7 (Paixão-Cavalcante et al., 2015). As well as their haemotoxic effects, these proteases are likely to be responsible for initiating the processes that lead to local tissue damage following puff adder bites, including swelling, blistering, inflammation and necrosis, the combined effects of which often result in severe pain. However, there have been no detailed studies of the puff adder proteins that might be responsible for this local pathology.

The abundance and variety of SVMPs and SVSPs in *B. arietans* venom has been confirmed by proteome analysis (Casewell et al., 2014; Fasoli et al., 2010). Along with the various studies on individual proteins, these proteomic studies have provided some insight into the toxicity of puff adder venom, but a complete picture of the protease complement, and its regional variations, is required to better understand the diverse pathological effects of the venom and identify therapeutic targets. The SVSP content of puff adders will be the subject of a separate study. Here, we describe the purification of, and the structural and functional characterisation of SVMPs that we have isolated from a variety of puff adder venoms. The study was prompted by our observation of differences in content and bioactivities of venom from Nigerian and Tanzanian snakes (Dawson et al 2024) and proteases isolated from these two form the bulk of this work, since transcriptomes are available for both. We have since widened the scope of this work, however, to determine the presence of the SVMPs in venoms from a regional variety of captive bred and wild-caught African and Arabian puff adders.

## 2. Materials and methods

### 2.1. Venom

*B. arietans* venoms were extracted from Nigerian (five snakes, named NGA B below),Tanzanian (four) and Eswatini (two) specimens presently maintained within the herpetarium at the Centre for Snakebite Research and Interventions at the Liverpool School of Tropical Medicine (LSTM). Five Kenyan *B. arietans* venoms were provided by the Kenyan Snakebite Research and Intervention Centre, Institute of Primate Research, Nairobi, Kenya. Venoms from individual specimens were labelled, immediately frozen at -20°C and later lyophilised for long term storage at 2 – 8°C. To deliver the volume of venom required for the multiple analyses described below, venom extractions were performed on multiple occasions from each snake. The other venoms used in this study were taken from stocks held at LSTM; these are either from snakes previously kept in the herpetarium (Ghana and Nigerian, the latter named NGA A herein) or samples taken from wild-caught animals over many years and supplied to LSTM. Immediately prior to biochemical analysis, the venoms were reconstituted in PBS (50 mM sodium phosphate, 0.15 M NaCl pH 7.2). For the chromatographic separations, the venoms were resuspended sodium phosphate buffers at the pH relevant to the method used for the first step in the procedure (see section 2.3).

### 2.2. Reagents

All reagents used were of analytical reagent grade and, unless otherwise specified, were purchased from Merck Life Science, Watford, UK or Fisher Scientific, Loughborough, UK.

### 2.3. Protein purification

All chromatography was carried out using either an AKTA LC system (Cytiva) or a Vanquish HPLC system (Thermo Fisher). Chromatography buffers were freshly prepared and vacuum filtered (0.2 um) immediately prior to use.

#### 2.3.1. Size exclusion chromatography of whole venom

The initial step in all isolation procedures was a separation of whole venom on a size exclusion chromatography (SEC) column. The buffers used for the SEC separation were chosen to facilitate the transfer to the second chromatography step (cation or anion exchange) without the need for buffer exchange. Thus, for SVMP PI isolation, 20 mg of freeze-dried venom was resuspended in 1.5 mL ice-cold PB5.2 (50 mM sodium phosphate pH 5.2) and centrifuged at 10,000 xg for 10 mins. The supernatant was immediately loaded onto a 120 mL column of Superdex 200HR equilibrated in PB5.2. For SVMP PIII isolation, all steps were identical but the PB5.2 buffer was replaced with 50 mM Tris-Cl, pH 8.5 (TB8.5). In both cases the column was operated at a flow rate of 1.0 mL/min and 2 mL fractions were collected after the void volume. Elution was monitored at 214 and 280 nm. SDS-PAGE analysis and protease assays were carried out on all protein-containing fractions to determine which to select for the second stage of isolation.

#### 2.3.2. SVMP PI purification

Using general protease assays, the bulk of the EDTA-inhibited proteolytic activities were found in the middle-eluting peaks of the SEC separations associated with prominent bands on SDS-PAGE at 20-30 kDa. These SVMP-containing fractions (2-3 mg of total protein) were applied to either a 1 mL HiRes Capto S or a 4.7 mL Capto S column equilibrated in PB5.2. Elution was carried using a 25-column volume (CV) gradient of 0 - 0.25 M NaCl in PB5.2. The flow rate was 0.6 – 1.0 mL/min. The unbound material was retained and 1 mL fractions were collected from the start of the NaCl gradient. Elution was monitored at 214 and 280 nm. Protease assays were carried out on the main peaks and SDS-PAGE and RP-HPLC was used to determine purity.

#### 2.3.3. SVMP PIII purification

The SVMPIIIs, including the prominent 68 kDa protein visible in some Kenyan *B. arietans* venoms were found in the first peak following the SEC separation of the venoms in TB8.5. The main SEC fractions containing the 50-70 kDa proteins visible on SDS-PAGE were applied directly to a 1 mL Mono Q column equilibrated in the same buffer. Bound proteins were eluted from the column using a 20 CV gradient of 0-0.3 M NaCl in TB8.5. The column was operated at 0.6 mL/min and 1.0 mL fractions were collected. Elution was monitored at 214 and 280 nm.

#### 2.3.4. Analytical methods

##### 2.3.4.1 SDS-PAGE

Samples were prepared for reducing SDS-PAGE analysis by adding sample buffer to a final concentration of 2% SDS, 5% β-mercaptoethanol and then heating at 85°C for 5 mins. Electrophoresis was performed on 4-20% acrylamide gels (BioRad TGX) using a Tris-glycine buffer system, followed by staining with Coomassie Blue R250, colloidal coomassie (ThermoFisher), silver-staining (Chevallet et al., 2006) or visualisation using a stain-free system (BioRad). Non-reducing gels were performed by omitting the heat treatment. The SVMP PIII isolated from Kenyan *B. arietans* venom required a different method of preparation for non-reducing gels in order to retain its oligomeric structure: thus, NP-40 was added to a final concentration of 0.5% before adding sample buffer containing 0.5% (final) SDS. Molecular weights of purified proteins were calculated using a calibration curve prepared from PageRuler markers (Thermo Fisher) run on the same gel (4-20% TGX, BioRad).

##### 2.3.4.2 Analytical SEC

For native size analysis of the Kenyan SVMP PIII, a 24 mL Superdex 200 SEC column was set up on an AKTA LC system (Cytiva) and equilibrated in PBS (25 mM sodium phosphate, 0.15 M NaCl, pH 7.2). The column was operated at a flow rate of 0.5 mL/min and elution was monitored at 280 nm. Fifty μL of sample was loaded. The column was calibrated by running 50 μL of BioRad SEC standard under the same conditions.

#### 2.3.5. Deglycosylation of isolated proteins

In preparation for deglycosylation, a 20 μL sample was denatured by heating for 5 mins at 85°C following the addition of 2.0 μL 1% SDS and 0.5 μL 1 M DTT. After cooling to room temperature,1.7 μL of 10% NP-40 was added and then 0.5 μL PNGase F (at the supplied concentration) was added to start the reaction. Incubation was carried out at 37°C for 2 hours. Where deglycosylation was carried out under native conditions for gelatin zymograms, the denaturing step was omitted.

#### 2.3.6. General protease assays

To distinguish between metalloprotease and serine protease activities, the ability of the purified proteases to degrade casein and insulin B was determined in the presence of EDTA (metalloprotease inhibitor). Insulin B degradation can also give an indication of the degree of specificity of action of the protease as evidenced by the number of peptide products observed on HPLC following digestion. Key SVMPs were also tested against the fluorogenic substrate ES010 since this is routinely used in our laboratory for high-throughput work.

##### 2.3.6.1 Casein degradation with SDS-PAGE analysis

Venom or protein samples were incubated in TBSC (25 mM Tris-Cl, 0.5 M NaCl, 1 mM CaCl_2_, pH 7.4) with beta casein at a ratio of 30:1 (w/w) casein:sample protein; using a final concentration of 1.0 mg/mL casein. Incubation was carried out at 37°C for 2 hours. SDS-PAGE sample buffer was added (final concentration 2% SDS, 5% β-mercaptoethanol) to stop the reaction. This was then heated at 85°C for 5 mins. The extent of casein degradation was assessed using SDS-PAGE (see 2.3.4.1). In EDTA inhibition experiments, EDTA was added to the protein sample to a final concentration of 5 mM and incubated for 15 mins prior to the addition of casein to start the reaction.

##### 2.3.6.2. Insulin B digestion with RP-HPLC analysis

Venom or protein samples were incubated in TBSC with insulin B chain at a ratio of 30:1 (w/w) insulin B:sample protein, using a final concentration of 0.02 mg/mL insulin B. Where EDTA was used, this were pre-incubated with the protein sample as in section 2.3.6.1. Incubation was carried out at 37°C for 90 mins and the reaction was stopped by the addition of trifluoracetic acid (TFA) to 1%. An aliquot containing the equivalent of 1 μg insulin B was analysed by RP-HPLC using a Biobasic C4 column (2.1 x 150 mm). The flow rate was 0.3 mL/min and the separation was carried with the following gradient of acetonitrile in 0.1% trifluoroacetic acid; 0-36%/40 mins; 36-70%/3 mins. Elution was monitored at 214 nm.

##### 2.3.6.4 Fluorogenic substrate assay

This assay, which kinetically measures the cleavage of a quenched fluorogenic substrate (ES010, R&D Biosystems) by metalloproteases was performed as previously described (Albulescu et al., 2020). Reactions were set up in a 384-well plate (Greiner) and consisted of 15 μL of pure SVMP PII isoforms at 0.05 – 0.2 mg/mL in PBS, and 75 μL of substrate (final concentration 10 μM) and were run in triplicate. Data was collected on a Clariostar (BMG Labtech) instrument at an excitation wavelength of 320 nm and emission wavelength of 405 nm at 25 °C for 1 h. The slope of the reaction between 0-2 min was calculated for each sample, the average background slope (PBS-only samples) was then subtracted, and specific activity expressed as ΔFluorescence/ time(min) /pmole protein.

#### 2.3.7. Functional protease assays

Five assays were carried out to determine the abilities of the isolated proteases to degrade proteins that may be of functional and pathological significance. Degradation of gelatin [predominantly collagen I] or laminin-rich basement membrane material are possible indicators of the ability to cause tissue damage. Potential coagulopathic activities were measured by the ability to degrade the key coagulation factors, fibrinogen, prothrombin and factor X. For the latter two substrates, specific proteolytic action required to generate the active forms (thrombin and factor Xa) can be determined.

##### 2.3.7.1. Gelatin zymogram assay

The gels (10% acrylamide) for this assay were prepared using the method of Laemlli (1970) with the adaptations of Fling and Gregerson (1986). Gelatin was prepared by briefly heating a 20 mg/mL solution at 50°C, then adding this to the gel mix immediately prior to casting, such that the final gelatin concentration was 2 mg/mL (0.2% w/w). Where inhibitors were tested, these were pre-incubated with the protein sample as in section 2.3.7.1. Venom or protein samples were then prepared for the zymogram assay by adding standard SDS-PAGE sample buffer, but with no reductant and a final SDS concentration of 1.6% (w/v). These were not heated. Molecular weight markers were run under standard reducing conditions, but with an empty lane between them and the sample lanes. Following electrophoresis, the method of Toth et al., (2012) was used to visualise gelatinase activity. Thus, the gel was rinsed, with shaking, in 2.5% (v/v) Triton X-100 for 30 mins, then washed (3 x 10 mins) with distilled water to remove the detergent. The buffer used for development was 50 mM Tris-Cl, 200 mM NaCl, 5 mM CaCl_2_, 0.02 % (v/v) Brij 100, pH 7.8. The gel was washed for 10 mins in this buffer, then this was replaced with fresh development buffer and incubated overnight at 37°C. To visualise gelatin degradation zones, the gel was stained for 60 mins with Coomassie Blue R250 and destained until clear bands were visible against a blue background.

##### 2.3.7.2. Digestion of basement membrane proteins

Basement membrane material (Geltrex, Thermo Fisher) was diluted with PBS to 10% of its supplied concentration (10-18 mg/mL) and stored in aliquots at -20°C. For the assay, a fresh aliquot was carefully thawed out and immediately placed on ice. Digestions were set up with basement membrane material:protein sample at 30:1 (w/w) and performed at 37°C for various time periods between 2 mins and 4 hours. Following this, SDS-PAGE sample buffer was added and the sample heated at 85°C to stop the reaction. The extent of basement membrane protein degradation was then assessed using SDS-PAGE (section 2.3.4.1).

##### 2.3.7.3. Digestion of plasma proteins (prothrombin, fibrinogen and Factor X)

In both the prothrombin and fibrinogen degradation assays, these two substrate proteins were used in the assay at a final concentration of 1.0 mg/mL in TBSC and venom/pure protein samples were added at a ratio of 30:1 substrate:protein sample to start the reaction. Incubation was carried out at 37°C for 60 mins. For the factor X activation, the final concentrations in the assay were 50 μg/mL factor X and 2.0 (3.5 for PIII) μg/mL purified SVMP. RVV-X was used a positive control in this assay: this was purified from Sri Lankan *D. russelli* according to the methods of Khin et al. (2017). This assay was carried out for 120 mins at 37°C. In all cases, EDTA was added (final concentration 5 mM) and incubated for 5 mins. to stop the reaction. SDS-PAGE sample buffer was then added and the sample was heated at 85°C for 5 mins. The extent of substrate protein degradation was then assessed using SDS-PAGE (section 2.3.4.1).

#### 2.3.8. Trypsin digestion and MS/MS analysis

In preparation for this, the relevant proteins were desalted on a RP-HPLC column, as above. These were then dried in a centrifugal evaporator, 20 μL H_2_O was added and then re-dried. These proteins were resuspended in 8 M urea/0.1 M Tris-Cl (pH 8.5), reduced with 5 mM TCEP (Tris (2-carboxyethyl) phosphine) for 20 minutes and alkylated with 50 mM 2-chloroacetamide for 15 minutes in the dark at room temperature. Samples were diluted 4-fold with 100 mM Tris-Cl (pH 8.5) and digested with trypsin at an enzyme/substrate ratio of 1:20 overnight at 37°C. The reaction was terminated by addition of formic acid (FA), and digested peptides were loaded on to Evotips and analysed directly using an Evosep One liquid chromatography system (Evosep Biosystems, Denmark) coupled with timsTOF SCP-mass spectrometer (Bruker, Germany). Peptides were separated on a 75 µm i.d. × 15 cm separation column packed with 1.9 µm C18 beads (Evosep Biosystems, Denmark) and over a predetermined 44-minute gradient. Buffer A was 0.1% FA in water and buffer B was 0.1% FA in acetonitrile. Instrument control and data acquisition were performed using Compass Hystar (version 6.0) with the timsTOF SCP operating in data-dependent acquisition mode.

Fragmentation spectra were searched against an in-house *B. arietans* venom gland derived protein sequence database sourced from Nigerian and Tanzanian specimens (see Dawson et al. 2024) using Mascot (Perkins et al., 1999). Reverse decoys and contaminants were included in the search database. Cysteine carbamidomethylation was selected as a fixed modification, oxidation of methionine was selected as a variable modification. The precursor-ion mass tolerance and fragment-ion mass tolerance were set at 10 ppm and 0.04 Da, respectively, and up to 2 missed tryptic cleavages were allowed. Mascot files were parsed into Scaffold (version 5.0.1, Proteome Software, Inc.) for validation at a protein-level false discovery rate (FDR) of < 1%.

## 3. Results

### 3.1. Comparison of the regional B. arietans venoms

Electrophoretic analysis of Nigerian (NGA B) and Tanzanian (TZA) venoms from individual snakes held at LSTM, Kenyan (KEN) venoms supplied by K-SRIC, and Ghana (GHA) venoms from stocks held at LSTM showed differences in their protein profile, but particularly so the prominent bands in an otherwise clear 20 – 30 kDa region on the gel (Fig. 1). TZA and GHA venoms possessed a sharp band at 21 kDa, whereas KEN and NGA B venoms had, instead, bands at 26 and 30 kDa respectively. In the higher molecular weight region of the gel, there was a lower level of the multiple protein bands in the 50-60 kDa SVMP PIII range compared with other African vipers such as Echis species. This region was particularly faint in the TZA venoms. One exception to this was a very prominent protein at 68 kDa that was seen in some of the KEN venoms (visible here in lane 3, snake 015), including one captive-bred KEN snake held at LSTM (not shown). These patterns were consistently observed in venoms from a wider selection of animals (Fig. S1). Interestingly two stock Nigerian venoms from snakes previously housed at LSTM (NGA A) contained this protein rather than the 30 kDa band observed in the venom of Nigerian snakes (NGA B) presently held at LSTM. Within the other venoms analysed for Fig. S1, a Saudi Arabian, a South African and three Namibian venoms contained the 30 kDa form. One of the Namibian venoms possesses a noticeably large amount of this protein (Fig S1, lane 3). This gel also emphasises the very low abundance in puff adder venoms of protein bands in the 50-65 kDa SVMP PIII range, though the Namibian venoms do appear to have more than can be seen in others. Suspecting that the prominent 21-30 kDa and 68 kDa protein bands were SVMPs, we set out to isolate them and test their activities towards key substrates.

**Fig. 1.**
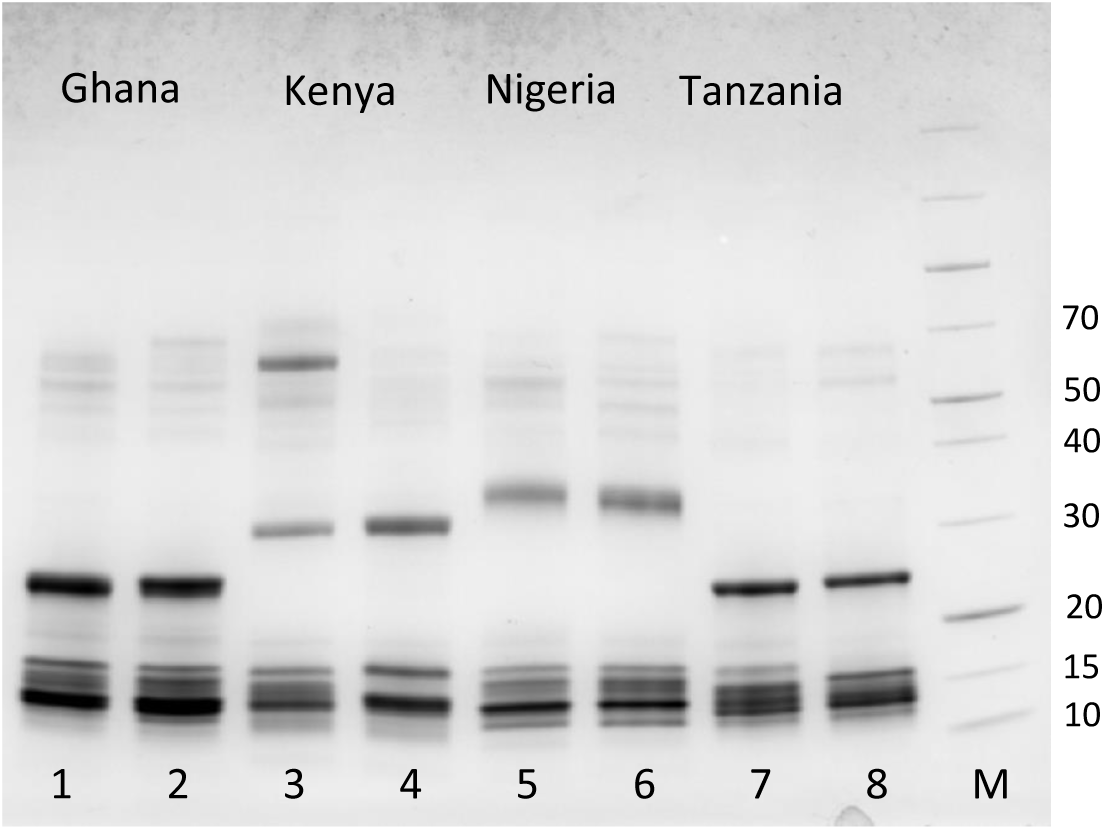
SDS-PAGE analysis of the venoms from individual *B. arietans* snakes from West and East Africa. The gel is 4-20% acrylamide (BioRad) and was run under reducing conditions. Eight μg of whole venom was loaded per lane and the gel and was stained using Coomassie Blue R250. The molecular weights of the markers (lane M, Thermo Page Ruler) are indicated in kDa. The venoms shown, two samples of each, are 1, 2: GHA; 3, 4: KEN; 5, 6: NGA B; 7, 8: TZA.

### 3.2. Isolation and biochemical characterisation of the SVMPs

The chromatography steps described here were performed multiple times, each time using venom from one individual snake or one batch of stock venom. The examples used here and in the supplement are for 20-30 kDa SVMPs isolated from Tanzanian, Eswatini and Nigerian (B) venoms obtained from LSTM-held snakes, from Kenyan venom provided by KSRIC, and from Ghanian, Nigerian (A) and Saudi Arabian stock venoms. Due to their low abundance, only small quantities of SVMP PIIIs were isolated from some of the venoms and, therefore, only the prominent 68 kDa protein in the Kenyan venoms is described in detail here.

#### 3.2.1. Isolation of the low molecular weight SVMPs

The suspected small SVMPs observed at 20-30 kDa on SDS PAGE of the various venoms were typically purified using SEC followed by cation exchange chromatography. An example trace for the SEC separation of the 21 kDa from Tanzanian (TZA) venoms can be seen in Fig. S2. The 21 kDa band, as seen in Fig 1, lanes 7 and 8, was found in the last of the 5 protein-containing peaks, 1-5. Further analysis (results not shown) revealed that peak 7 is adenosine and peak 6 is mostly the tripeptide SVMP inhibitor. The 21 kDa protein was fairly pure at this stage but was further purified using cation exchange chromatography (Fig. S3). The protein eluted in two peaks, the second of which contained the greater amount of the protein.

Fig S4 shows a typical SEC separation of a Nigerian (NGA B) venom containing the 30 kDa protein, quite different in profile to that of the TZA venoms. The 30 kDa NGA (B) protein was found in the first half of the split peak number 3 and so the proteins in these fractions were subject to cation exchange chromatography (Fig 5S). The 30 kDa protein was found to be pure in both of the prominent peaks eluting at 11-17 mL and the irregular peak shapes suggest it exists in multiple forms.

Using the insulin B and casein assays carried out in the presence or absence of EDTA, both proteins were confirmed to be SVMPs (Fig 7S and 8S). Although both the 30 kDa (NGA B) and 21 kDa (TZA) SVMPs were able to fully digest insulin B chain within 90 mins, the patterns of digestion for the two are different suggesting differences in their peptide bond specificities. The 30 kDa SVMP appears to have cleaved at just two sites, generating three products, whereas 6-7 peptides have been produced by the 21 kDa SVMP. As will be seen in subsequent assays, the 21 kDa SVMP is clearly a more a potent protease than the 30 kDa SVMP.

Because the 26 kDa protein seen in the Kenyan (KEN) venoms has a similar native molecular weight to the abundant multiple forms of CLP dimers found in puff adders, SEC wasn’t effective as part of the isolation procedure. It was therefore isolated using a double cation exchange protocol (result for second step on Fig. S6). This was also confirmed to be a metalloprotease using casein and insulin B degradation assays (Fig. S7 and S8). The two-peptide insulin B degradation pattern suggests a narrow specificity, cleaving at just one position, one that appears to overlap with the 30 kDa (NGA B) SVMP based on the co-elution of its two peptide products with two of the three in the latter.

21 kDa (TZA) and the 30 kDa (NGA B) SVMPs were subjected to de-glycosylation with PNGase F (Fig. 9S). The 21 kDa (TZA) SVMP appears to be non-glycosylated as there was no change in mobility on SDS-PAGE following PNGase F treatment. In contrast, the 30 kDa (NGA B) SVMP shifted in molecular weight by ∼9 kDa, which suggest the presence of 3 N-glycans (assuming 2.5-3 kDa per glycan), but a time-course de-glycosylation indicated only one intermediate (result not shown) and that it therefore possesses 2 N-glycans. De-glycosylation of the 26 kDa (KEN) SVMP caused a mobility shift on SDS-PAGE of ∼5 kDa (Fig. S6 inset B) and, taking into account the results for the 30 kDa form, is most likely to possess just one N-glycan. Following de-glycosylation, all three proteins ran at the same molecular weight (∼21 kDa). When PNGase F treatment of the 26 and 30 kDa SVMPs was carried out under native conditions (in the absence of DTT, SDS and NP40), the proteins became completely insoluble (result not shown).

MS/MS analysis of tryptic peptides derived from 21 kDa (TZA) and 30 kDa (NGA B) proteins determined that they are both SVMP PIs, derived from an SVMP PII transcript. The two sequences to which these matched, found in transcriptomes from the respective snakes, are shown in Fig. 2. The transcripts for these two proteins were top of their respective list (all proteins) for transcripts per million (Dawson et al. 2024), reflecting the prominence of the proteins in the venom as visualised by SDS-PAGE (Fig. 1). No disintegrin-derived peptides were found in the MS/MS analysis. The two proteins match with the predicted molecular weight of the metalloprotease domain of the SVMP PIs, and the number of predicted N-glycan sites for each protein matched to that found experimentally. No transcript information was available for KEN *B. arietans* at the time of writing, so it was not possible to match the 26 kDa (KEN) SVMP to a specific SVMP sequence.

**Fig. 2.**
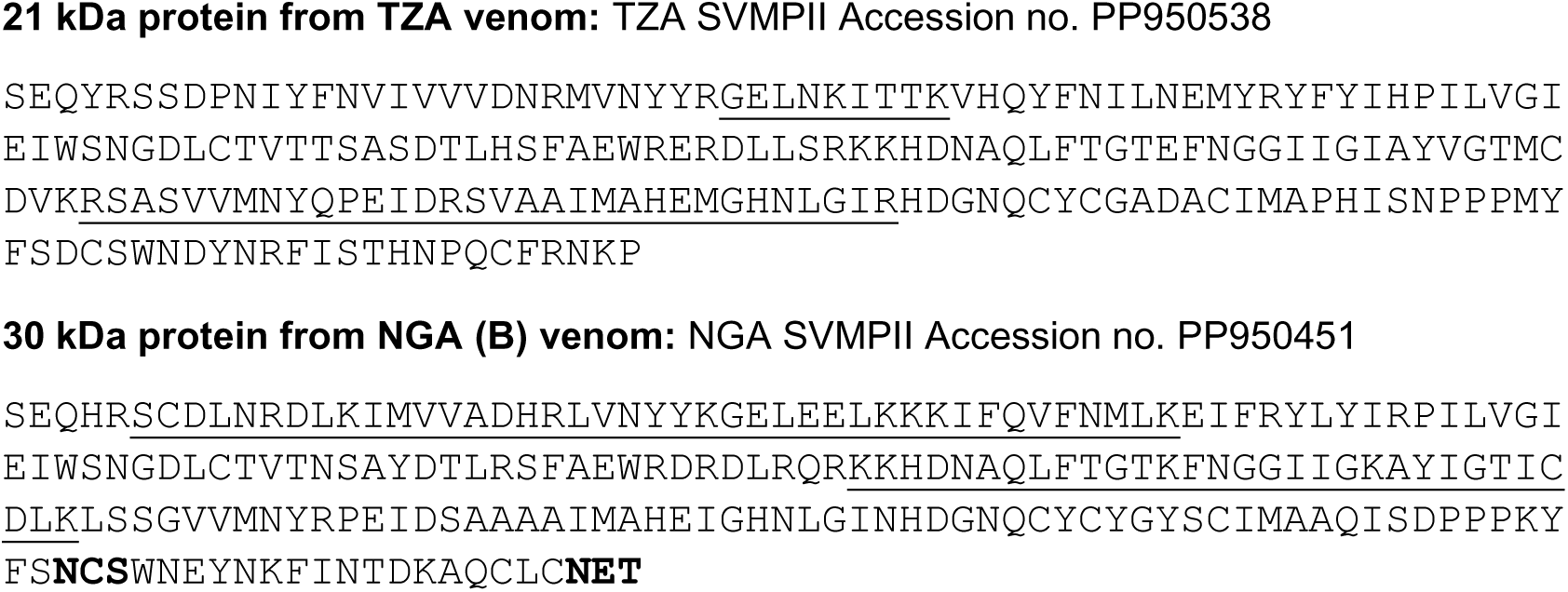
Amino acid sequences of the transcripts matching to the SVMPs isolated from TZA and NGA (B) venoms. Ten μg of the pure protein was digested with trypsin and the resulting peptide mix was analysed by LC-MS/MS (Sec. 2.3.8). Fragmentation spectra were searched against an in-house *B. arietans* (NGA and TZA) venom gland derived protein sequence database (Dawson et al. 2024) using Mascot. The tryptic peptides identified by MS/MS analysis are underlined and the predicted N-glycan sites in the NGA sequence are indicated in bold font.

Since these SVMPs are likely to be therapeutic targets for managing envenoming by *B. arietans*, the activity of 21 kDa (TZA) and 30 kDa (NGA B) SVMP PIs was measured against a fluorogenic peptide MMP substrate routinely used in our laboratory for high-throughput drug discovery work. The 21 kDa (TZA) SVMP PI was found to have a much greater specific activity than the 30 kDa (NGA B) form under the conditions used: 2290 +/- 211 units (ΔFluorescence/min/pmole) compared with 148 +/- 1.2 units respectively. This pronounced difference in protease activity is mirrored in the functional assays carried out on the proteins (see Sec. 3.3).

Using the same isolation methods as described above, 21 and 30 kDa forms of the SVMPs were also isolated from some of the venoms used for the SDS-PAGE analysis in Fig. S1. A 21 kDa form was isolated from stock Ghana (GHA) venom (Fig. 1S10) and a 21 kDa form was also isolated using just SEC of venom from an Eswatini (ESW) snake kept at LSTM (Fig. 11S). Stock venoms from Nigerian snakes previously housed at LSTM (NGA A) all contained a 21 kDa form (see Fig. S1, lanes, 1-2) and this was isolated from one of these (Fig. S12). A 30 kDa form was purified from a Saudi Arabian (SDA) venom stock (Fig. S13). The latter was subjected to deglycosylation with PNGase F resulting in a 9-10 kDa decrease in molecular weight (Fig. S13 inset B), suggesting the presence of 2 N-glycans as in the case the 30 kDa NGA (B) SVMP PI.

#### 3.2.2. Isolation of the high molecular weight SVMP from Kenyan venom

SEC of the KEN (snake 015) venom resulted in a large peak eluting at the start of the separation (Fig. S14). This contained the 68 kDa protein visible on SDS-PAGE of whole venom (see Fig. 1, lane 3). The fractions containing this protein were subject to anion exchange chromatography resulting in two peaks (Fig. S15), the first of which contained the 68 kDa protein at a high degree of purity (Inset Fig. 15S and Fig. 3, lane 1). The second peak contained a protein with the α/b subunit pattern expected of a CLP, presumably a high molecular weight form (e.g. tetrameric) as commonly found in many other vipers (Arlinghaus and Eble, 2012). The 68 kDa protein was determined to be an SVMP using a casein assay +/- EDTA (Fig. S8); it is presumably an SVMP PIII because of its molecular weight. Unlike the SVMP PI forms, this SVMP PIII was unable to cleave any bonds in insulin B (Fig S7). Because of its elution position on SEC and suspecting it to be responsible for the 140 kDa gelatinase activity in the KEN015 venom (see below), it was expected to be a dimer (an SVMP PIII-c) but upon analysis using SDS-PAGE under non-reducing conditions (Fig. 3) the protein still ran at ∼68 kDa, even when using a low SDS concentration (0.5% w/w final). If, however, the protein was treated with the non-ionic detergent NP-40 prior to the addition of a low % SDS non-reducing sample buffer, then it remained native and ran at 140 kDa (Fig. 3, lane 4). Its native molecular weight was also confirmed using analytical SEC, where it eluted with a molecular weight of 170 kDa, though a small proportion has clearly broken down into monomeric form, eluting at 70 kDa (Fig. 4).

**Fig 3.**
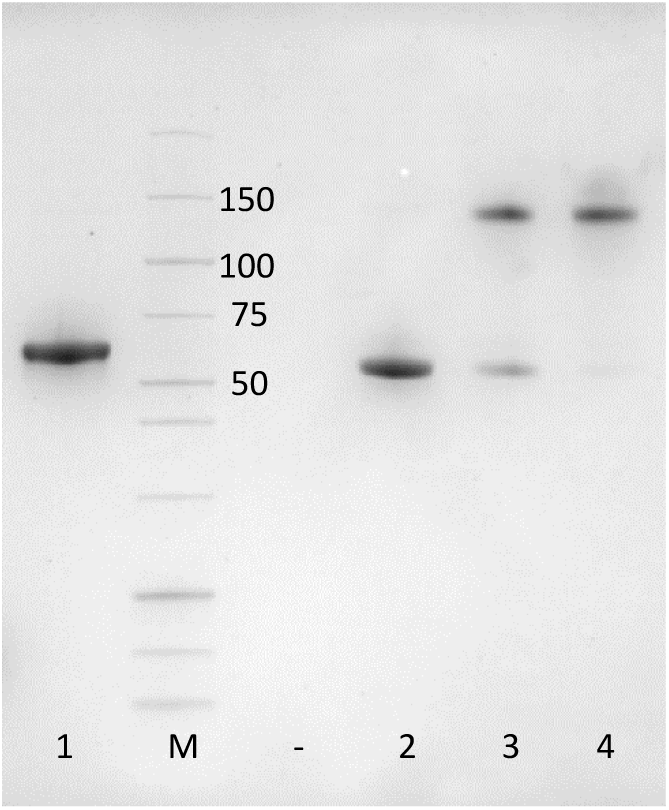
Effect of non-ionic detergent on the oligomeric structure of the KEN SVMP PIII. SDS-PAGE of KEN SVMP PIII run under various conditions. Lane 1, standard reducing conditions; lane M, molecular weight markers (Thermo PageRuler with key molecular weights indicated in kDa), lane 2 non-reducing conditions (0.5% SDS), lane 3 as lane 2 but with 0.25% NP40 added after the SDS, lane 4, as lane 2 but with 0.25% NP40 added prior to the SDS. The gel used was a 4-20% acrylamide (BioRad) and was stained with Coomassie Blue R250.

**Fig. 4.**
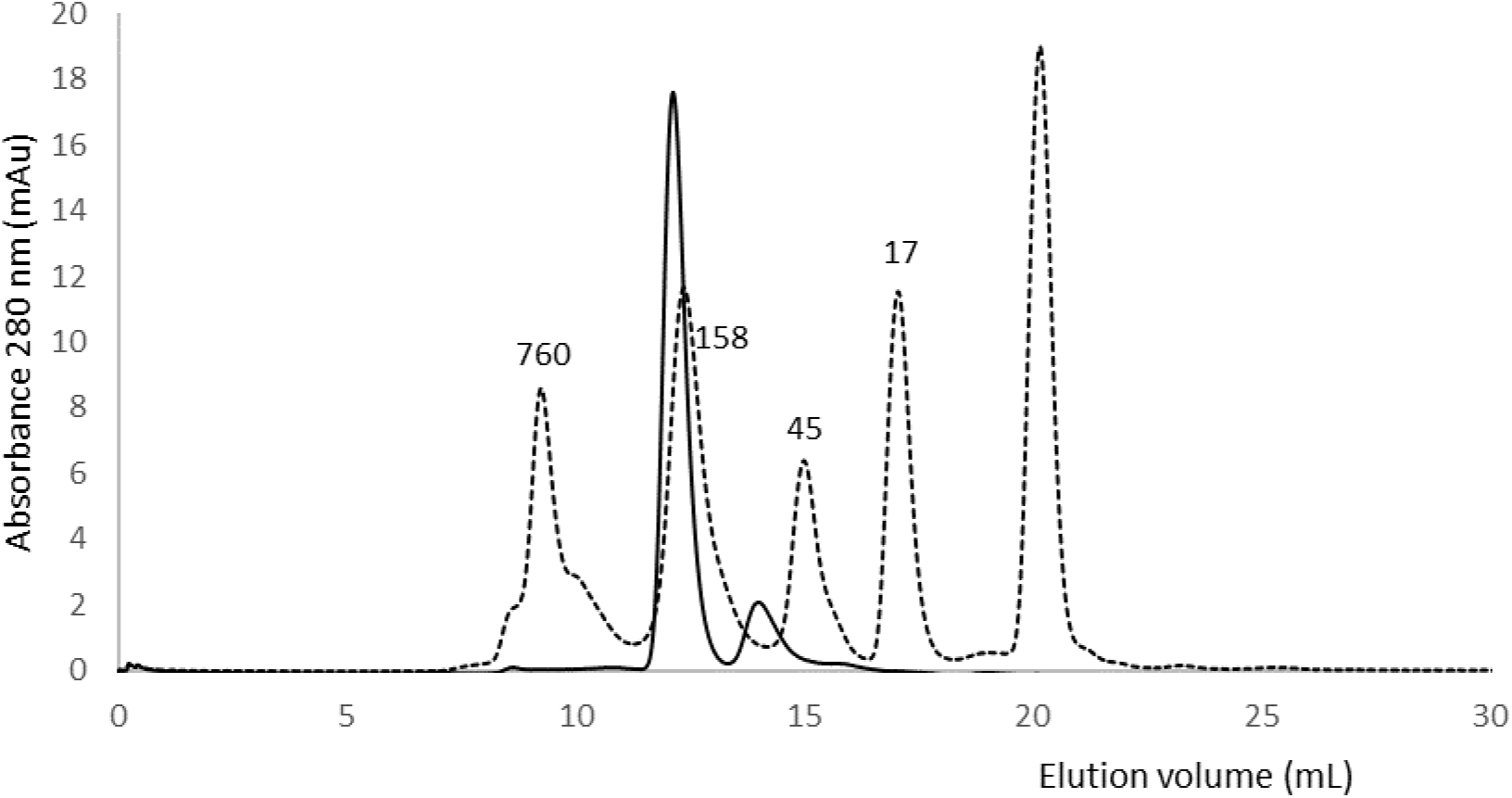
Size exclusion chromatography analysis of KEN SVMP PIII. Analysis was carried out using a 24 mL Superdex 200 SEC column operating at 0.5 mL/min in 25 mM sodium phosphate, 0.15 M NaCl, pH 7.2. Elution was monitored at 280 nm. Fifty μL (20 μg) of sample was loaded. Solid line, pure KEN SVMP PIII (peak 1 in Fig. 9S); dashed line, SEC standards (BioRad: ferritin, IgG, ovalbumin, myoglobin, cyanocobalamin) with native molecular weights (kDa) of the proteins indicated above the respective peak.

PNGase F treatment showed the KEN SVMP PIII was likely to possess 3 N-glycans, based on the 9-10 kDa shift in molecular weight that was observed (Fig. S9). This was confirmed by de-glycosylating under native conditions: bands indicating 1- and 2-glycan intermediates where observed (result not shown).

### 3.3. Activity of the isolated proteases in the functional assays

Preliminary to this work, we had found pronounced differences in the ability of venoms from the main snakes used here to degrade substrate proteins traditionally used to indicate coagulopathic and tissue-degrading activities of snake venoms. Those containing the 21 kDa SVMP PI (GHA, NGA A, ESW, TZA) were far more potent in their degradative action against fibrinogen, prothrombin, factor X and basement membrane proteins. On the other hand, all the NGA venoms (NGA A and B) used here, as well as those from ESW and GHA, possessed a strong 40-50 kDa gelatinase activity that was absent in all of the TZA and KEN venoms, and the one ZAF venom in our possession. The exception to the latter was a very active, high-molecular weight (∼ 140 kDa) gelatinase observed in some KEN venoms. Using the purified proteases, we were able to identify the factors responsible for these differences.

#### 3.3.1. Plasma proteins: Fibrinogen, prothrombin and factor X degradation

The results of these assays mirrored that of the biochemical assays; the 21 kDa SVMPs were far more potent than the 30 kDa glycosylated forms. All of the isolated SVMPs were able to digest fibrinogen to some degree (Fig. 5A). Complete degradation of both alpha and beta sub-units was observed when fibrinogen was incubated with each of the four 21 kDa forms (GHA, NGA A, ESW and TZA), and in the case of the GHA protein, there was even significant loss of the gamma subunit (lane 1). In contrast the two 30 kDa glycosylated forms (SDA and NGA B) appeared to have only alpha-fibrinogenase activity, with possibly a little beta subunit degradation by the NGA (B) form. The smaller 26 kDa glycosylated form from KEN venom (lane 5) was as active towards fibrinogen as some of the 21 kDa forms.

**Fig. 5.**
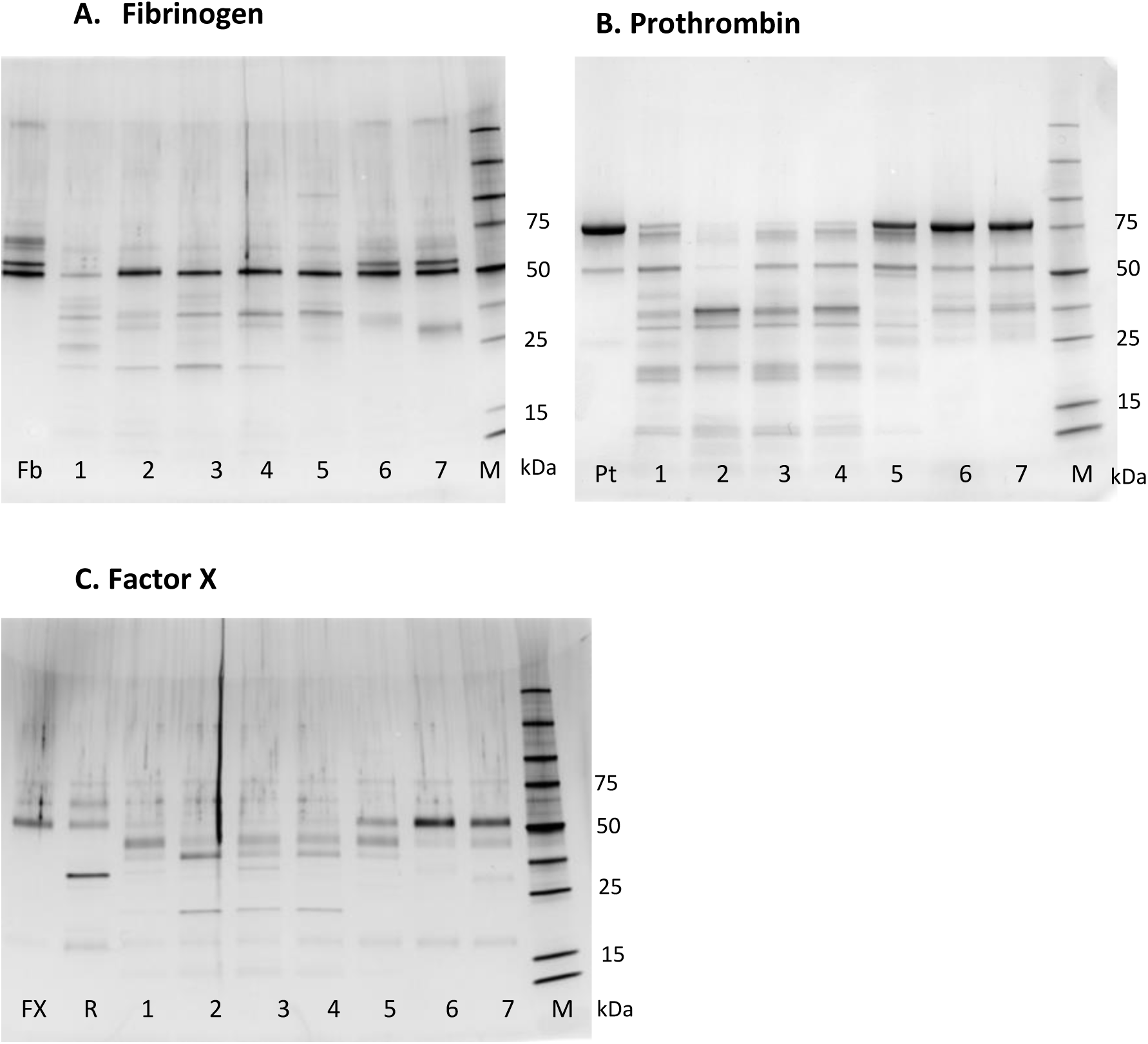
Degradation of fibrinogen, prothrombin and factor X by purified *B. arietans* SVMP PIs. Fibrinogen (Gel A), prothrombin (Gel B) or factor X (Gel C) were incubated at 37°C for 120 minutes with the proteases at a 30:1 (w/w) ratio of substrate:protease. The gel was run under reducing conditions and the equivalent of 1.0 μg (prothrombin, fibrinogen) or 0.4 ug (factor X) of substrate protein was loaded per lane. The gels are 4-20% acrylamide (BioRad) and were stained using Coomassie Blue R250 (prothrombin) or silver stain (fibrinogen and factor X). Lanes: Fb, Pt and FX, fibrinogen, prothrombin and factor X controls (no SVMP); R: RVV-X with factor X; lane 1, GHA 21 kDa SVMP; lane 2, NGA A 21 kDa SVMP; lane 3, ESW 21 kDa SVMP; lane 4, TZA 21 kDa SVMP PI; lane 5, KEN 26 kDa SVMP; lane 6, NGA B 30 kDa SVMP PI; lane 7, SDA 30 kDa SVMP. The molecular weights of key markers (lane M, Promega Broad Range) are indicated in kDa.

These relative levels of proteolytic activity were also reflected in the prothrombin degradation assay (Fig. 5B). Once again, the 21 kDa forms were the more destructive, though in this case the NGA (A) 21 kDa form appeared to possess the greater activity under the conditions used here (lane 2). This degradation is clearly non-specific and there is very little evidence of the B-thrombin (31 kDa) band that we routinely and clearly see with Echis sp. SVMPs. Within the three glycosylated forms (KEN, NGA B, SDA, lanes 5-7), the 26 kDa KEN form has been more active against prothrombin, as was the case with fibrinogen.

This pattern was again seen in the proteolytic action of the SVMPs against factor X (Fig. 5C). The 21 kDa forms caused more degradation than the 30 kDa NGA and SDA glycosylated forms, and the 26 kDa KEN form was intermediate in this respect. Critically, however, the action was non-specific and in no case was the 32 kDa truncated FXa H-chain (see RVV-X control, lane R) observed that is indicative of factor X activation.

The KEN 68 kDa SVMP PIII was tested against the same set of substrates (Fig. 6). This showed clear alpha and beta-fibrinogenase activity. As with the smaller SVMPs, the SVMP PIII was active against prothrombin, but this activity was weak and non-specific (no B-thrombin band was observed). There was no activity at all against factor X, even when the SVMP PIII was used at a higher enzyme:substrate ratio (∼1:15) than was used in other assays.

**Fig. 6.**
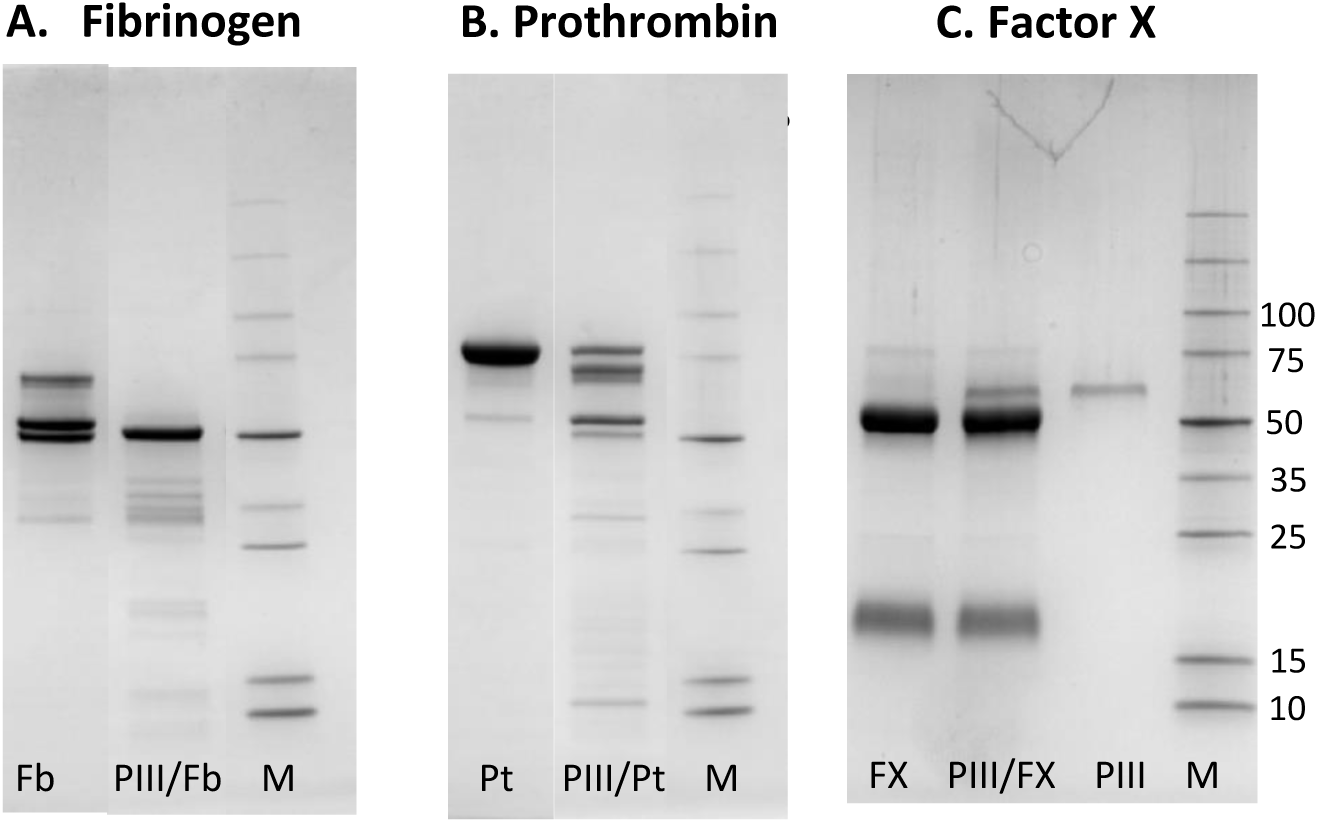
Degradation of fibrinogen and prothrombin by purified *B. arietans* SVMP PIII. Fibrinogen (Gel A) or prothrombin (Gel B) or factor X (Gel C) were incubated at 37°C for 120 minutes with the proteases at a 30:1 or 10:1 (factor X) ratio of substrate:protease. The gel was run under reducing conditions and the equivalent of 1.0 μg of substrate protein (prothrombin, fibrinogen, factor X) was loaded per lane. The gels are 4-20% acrylamide (BioRad) and were stained using Coomassie Blue. Lanes: Fb, Pt and FX: controls, no enzyme; PIII/Fb, PIII/Pt and PIII/FX: enzyme + relevant substrate; PIII: enzyme alone (factor X assay only). The molecular weights of key markers (lane M, Promega Broad Range) are indicated in kDa next to Gel C

#### 3.3.2. Basement membrane protein degradation

Using a similar assay method to that used for the plasma proteins, a basement membrane protein extract (Geltrex) was incubated with the purified 21-30 kDa SVMPs (Fig. 7A). Once again, the 21 kDa SVMPs were the most destructive, almost completely degrading both laminins under the conditions used here. The 30 kDa SDA and NGA (B) proteins appeared to have had no effect on either of the laminins. The 26 kDa KEN form was intermediate between the two, showing some degradation of the α laminin. The KEN SVMP PIII had little, if any, effect on the laminins (result not shown). The 21 kDa (TZA) and 30 kDa SVMP (NGA B) PIs were tested for their speed of action in a time-course experiment (Fig. 7B): significant levels of digestion by the 21 kDa SVMP PI were seen against laminin and the band at 130 kDa (most likely nidogen, Escalante et al., 2006) within 5 mins of incubation, whereas the 30 kDa (NGA B) SVMP PI had little effect, even after 30 mins.

**Fig. 7.**
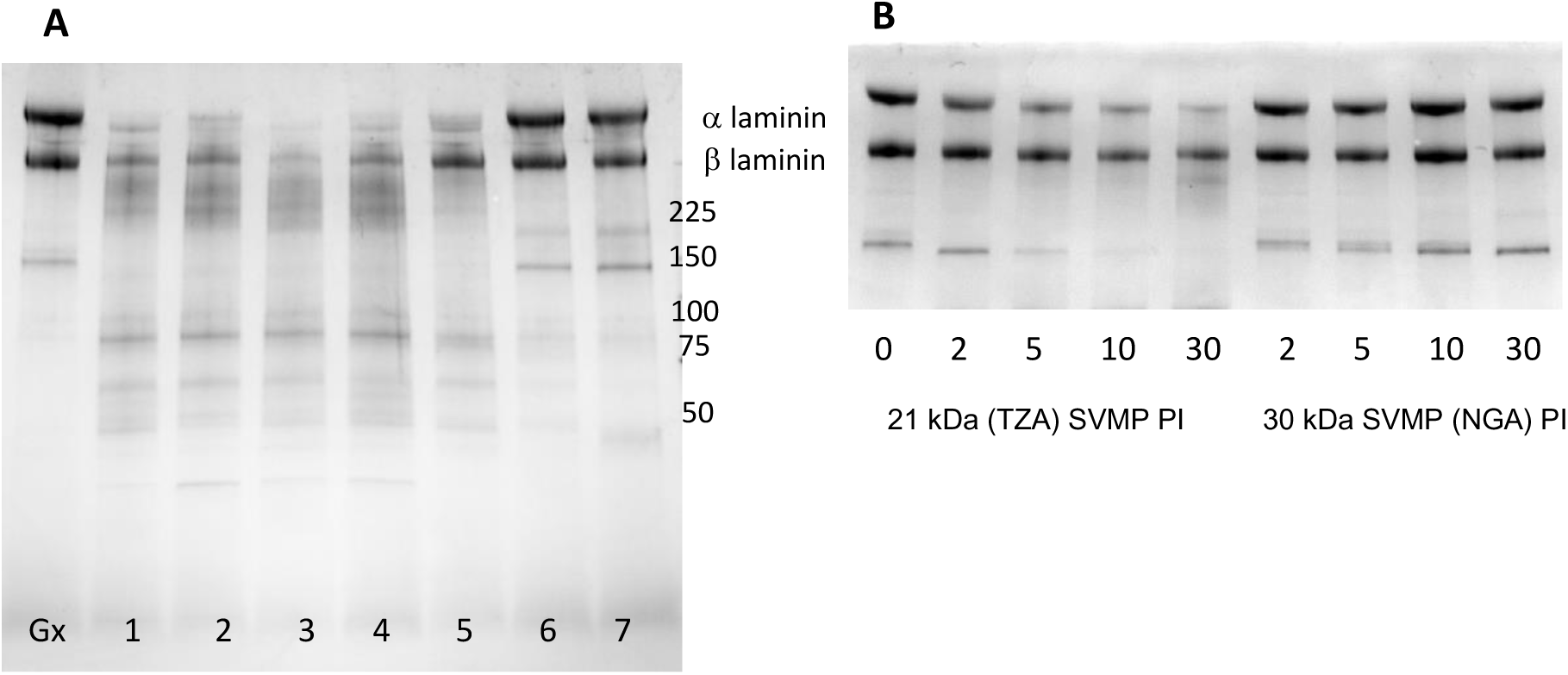
Degradation of basement membrane proteins by SVMPs isolated from *B. arietans* venoms. A basement membrane protein extract (Geltrex) was incubated at 37°C for 30 minutes with the purified proteases at a 30:1 (w/w) ratio of substrate:protease. The gel was run under reducing conditions and the equivalent of 8 μg of substrate protein was loaded per lane. The gel is 4-20% acrylamide (BioRad) and was stained using Coomassie Blue R250. **Gel A:** Activity of the individual SVMPs: Lanes: Gx, controls (no SVMP); lane 1, GHA 21 kDa SVMP PI; lane 2, NGA A 21 kDa SVMP PI; lane 3, ESW 21 kDa SVMP PI; lane 4 TZA 21 kDa SVMP PI; lane 5, KEN 26 kDa SVMP; lane 6, NGA B 30 kDa SVMP PI; lane 7, SDA 30 kDa SVMP. The molecular weights of key markers (lane M, Promega Broad Range) are indicated in kDa. **Gel B:** Time course of degrading activity of the 21 kDa (TZA) SVMP PI and 30 kDa SVMP (NGA B) PI; reaction was stopped after 2-, 5-, 10- and 30-mins incubation; lane 0, basement membrane protein control.

#### 3.3.3. Gelatinase activity

Contrasting levels of gelatinase activity were found in the puff adder venoms using a zymogram assay: in particular, some had a very strong activity in the 40-50 kDa region on the zymograms that was completely absent in others. Using inhibitors (EDTA and PMSF), this activity was shown to be due to a novel serine protease found only in some puff adder venoms (NGA A and B, GHA, SWZ and Malawi) and is the subject of a separate study on Bitis SVSPs (manuscript in preparation). None of the purified 21 – 30 kDa SVMPs showed any gelatinase activity. In the KEN venoms, however, a high-molecular weight (∼140 kDa), EDTA-inhibited gelatinase activity was identified that wasn’t present in any of other venoms used in this study. The dimeric SVMP PIII isolated from the KEN venom 015 (sec. 3.2.2) was shown to be responsible for this gelatinase activity (Fig. 8). Under the electrophoresis conditions used in this zymogram assay, NP40 pretreatment is needed to maintain the dimer form (as demonstrated in section 3.2.2). In lane 1, run in the absence of NP40, the clear band at 140 kDa is due to a small amount of remnant active dimer, but the bulk has broken down to an inactive monomer under these conditions, visible as a coomassie-stained band at around 55 kDa. When run after prior treatment with NP-40 (Fig 8, lane 2) however, all of the SVMP PIII remained in the fully active 140 kDa dimeric form. It is clearly only active in its dimer form. Its N-glycans are not essential for the gelatinase activity of the SVMP PIII. Thus, lane 3 shows the shift in molecular weight (Ca. 10 kDa) due to removal of the N-glycans by PNGase F, but the dimer form of the de-glycosylated protein is still able to degrade gelatin. Despite the presence of NP40, some of the de-glycosylated SVMP PIII has broken down to inactive monomer (stained band at around 45 kDa) following de-glycosylation, suggesting that the N-glycans may play some role in maintaining structural stability and/or solubility. (Note that the molecular weights of the PIII on the zymogram differ slightly to that on the standard gels in Figs. S9 and S15 due to the two different running conditions).

**Fig 8.**
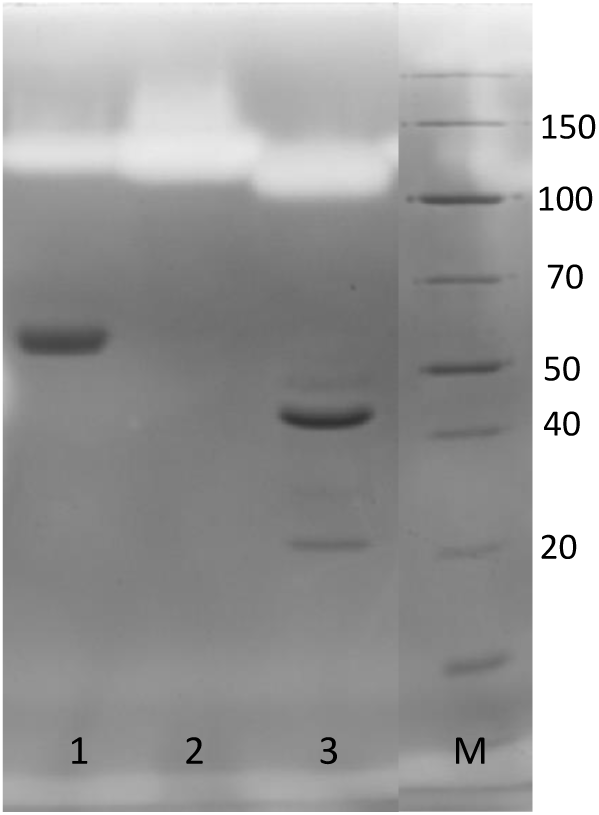
Gelatinase activity of native and de-glycosylated KEN SVMP PIII. The zymogram was prepared with 0.2% (w/w) gelatin in a 10% acrylamide SDS-PAGE gel. The gel was run under non-reducing conditions. The zymograms were then incubated as in section 2.3.7.1 and then visualised by staining with Coomassie Blue R250. This zymogram was destained for sufficient time to visualise the Coomassie-stained SVMP PIII bands. Lane 1, untreated SVMP PIII; lane 2 (control) and 3 (+ PNGase F), deglycosylation in the presence of 0.4% NP-40; lane M, molecular weight markers (Thermo PageRuler with molecular weights indicated in kDa).

## 4. Discussion

Our study on puff adder venoms has revealed the presence of an interesting set of SVMPs: an SVMPI, present in high abundance in either a glycosylated or a non-glycosylated form, and a gelatinolytic SVMP PIII with a novel dimeric structure. Measurements of their activities against major extracellular matrix proteins and key proteins of the coagulation pathway point to a likely role in the pathology of puff adder envenomation.

In the venoms studied here for which transcripts are available, TZA and NGA, we were able to demonstrate that the predominant form of SVMP is a PI which processed from an SVMP PII transcript, following the scheme of Fox and Serrano (2008). A BlastP search demonstrated these two SVMPs have little homology with other venom proteases, with no matches to non-*B. arietans* proteins scoring higher than 52%. As well as the clear visibility on SDS-PAGE of crude venoms, the abundance of these SVMP PI proteins within puff adder venoms is very evident in the proteomic analysis in Caswell et al. (2014, Supplement), though these proteins were not identified at the time, and in the transcriptome data of Dawson et al., (2024). It is not unreasonable to speculate that the equivalent proteins isolated from the KEN, GHA, ESW and SDA venoms and those observed on SDS-PAGE in other venoms are also of this class. Because of the predominance of these SVMPs in puff adder venom, and their undoubted pathological significance, we propose the name arilysin for these puff adder (*B. arietans*) SVMP PIs. Although the full SVMP PII protein (metalloprotease + disintegrin) and disintegrin domains derived from processing of these have often been isolated and structured, particularly from *Bothrops* and *Trimeresurus* spp., SVMP PI proteins from PII transcripts have not been identified in many other venoms (Olaoba et al., 2020). The most studied of these has probably been atrolysin E of *C. atrox* (Hite et al., 1992, Shimokawa et al., 1996) which, like the 21 kDa SVMP PI isolated here, has been shown to degrade extracellular matrix components (Baramova et al., 1989).

Of most interest amongst the arilysins is the very potent 21 kDa SVMP form found in TZA, GHA and SWZ venoms and also in some NGA venoms. This is a highly active protease with a broad specificity, acting strongly and rapidly against all the proteins used as substrates in this study. Based on its identical molecular weight and two-peak elution pattern on cation exchange chromatography (Fig. S3), this is almost certainly the broad specificity ‘Protease A’ isolated from puff adder venom (most likely South African) by Van Der Walt and Joubert (1971, 1972) and there are good matches between the sequence of 21 kDa SVMP PI and those of tryptic peptides from Protease A sequenced by this group in a subsequent paper (Strydom et al., 1986). What is particularly notable about the 21 kDa arilysin from TZA venom was the speed with which it acted against the proteins in the basement membrane extract (Fig. 7B), causing significant laminin degradation within minutes of exposure. One of the doubts over using this particular assay to determine potential haemorrhagic abilities of proteases are concerns on the duration of the assays and their clinical relevance, with the proteases showing their effects only after several hours in this in vitro assay (Escalante et al., 2006), much longer than the time taken to observe haemorrhage in a clinical setting. That is not the case for the 21 kDa arilysin, which acts within minutes.

The higher molecular weight of the three 26-30 kDa arilysins isolated from KEN, SDA and NGA venoms was shown experimentally to be due to N-glycosylation. The 30 kDa NGA form has two N-glycans (these sites were also identified in the primary structure, see Fig. 2) as does 30 kDa SDA form. The smaller 26 kDa KEN SVMP appears to have only one N-glycan, however. As measured by SDS-PAGE, all three forms were the same molecular weight as the 21 kDa forms of the protein following de-glycosylation. Though not unique to these puff adder SVMPs, glycosylated SVMP PIs are unusual. Rhodostoxin, an SVMPI found in *C. rhodosotoma* (Chung et al., 1996) and which is also derived from an SVMPII transcript, was found to have two N-glycans, one of which is at an equivalent site to one of those in the 30 kDa (NGA B) SVMP PI (see Fig. 2; PPPKYFS**NCS,** compared with PPSKYFS**NCS** in rhodostoxin). In a subsequent study, Tan et al. (1997) found that removal of the N-glycans from rhodostoxin did not affect its haemorrhagic activity. We were not able to measure the effect of N-glycan removal on protease activities because the proteins lost solubility as a consequence. However, the pronounced differences in activity - against all protein substrates - between the non-glycosylated 21 kDa arilysins and the glycosylated 30 kDa arilysins may be due to the presence of the two N-glycans, possibly hindering full access to larger substrates. The intermediate activity observed throughout for the 26 kDa KEN arilysin, which appears to have just one N-glycan, also supports this idea.

Since the specific activity towards the small ES010 substrate is much greater in the 21 kDa than in the 30 kDa arilysins, it is likely that the differences in activity between the two forms also resides in the primary structure, particularly at and around the active site. Interestingly, analysis of the venom gland transcriptomes (Dawson et al. 2024) showed regional differences in the tripeptide SVMP inhibitor (SVMPi) expressed by the puff adders (Wagstaff et al., 2008). The NGA venom from which the 30 kDa arilysin was derived has pELW (the presence of this in the venom has been confirmed in our laboratory) in the relevant transcriptome, whereas the TZA venom transcriptome predicts an SVMPi with the sequence pEVW. An aliphatic residue in the middle position of an SVMPi is unusual in itself, but, also, as far as we are aware, this is the first example of sequence variation of SVMP inhibitor tripeptides within the same species and might suggest that there are structural differences at the substrate-binding sites of the 21 and 30 kDa arilysins in which the SVMPi will reside in the venom gland.

Although they are not as visible on SDS-PAGE as they are in other viper venoms such as *Echis* spp., SVMP PIIIs at around 60-65 kDa were isolated from TZA, NGA, GHA and SWZ venoms (results not shown). These were found to be present only in very small quantities, however, and were therefore considered not to be of major clinical relevance. The low abundance of SVMP PIIIs in puff adders has also been identified in previous proteomics and transcriptomic studies (Caswell et.al. 2014, Juarez et al. 2009, Dawson et al. 2024). An SVMP PIII was found in the KEN venoms, however, that was particularly prominent in the venom of one animal, highly visible on SDS-PAGE of crude venom (Fig. 1, lane 3). Because of its abundance in the venom, this SVMP PIII was easily isolated and proved to be quite an interesting protein, having a narrower specificity than other viper SVMPs. It was not active at all against insulin B, a commonly used assay for venom SVMPs (Yamakawa et al., 1995). Although it showed some ability to degrade fibrinogen, as most venom proteases seem to do, it showed very little activity against basement membrane proteins and prothrombin and none against factor X. However, the purified protein did turn out to be responsible for the strong gelatin-degrading activity of the Kenyan venom. It is unlikely that this protein is confined to Kenyan snakes: A high molecular weight gelatinase activity was observed in 1 of 12 Nigerian *B. arietans* tested venoms in an earlier study (Currier et al., 2010) and this correlated with a pronounced ∼65 kDa band seen after western blotting with an SVMP antibody. Paixão-Cavalcante et al., (2015) also observed a high molecular weight gelatinase activity in a mix of puff adder venoms from Guinea, São Tomé, Angola and Mozambique. The variable presence of this SVMP PIII gelatinase in puff adder venoms suggests that is a protein that is only expressed under certain circumstances.

This Kenyan SVMP PIII possesses a novel oligomeric structure. It appears at around 65 kDa on reducing and non-reducing SDS-PAGE, but analytical SEC and non-reducing PAGE in the presence of NP40 showed that it exists naturally as a dimer. Gelatinase activity was observed only when the protein was maintained in this dimeric form. Each monomer was also found to possess 3 N-glycans which were shown to be unnecessary for its gelatinase activity but, as was the case for the SVMP PI, appear to be important in maintaining solubility. Interestingly, the dimer is clearly not held together through disulphide bonds, as would be the case it if were an SVMP of the PIII-c class and, as such, is a novel form of SVMP PIII, not presently accounted for in the SVMP classification system (Fox and Serrano, 2008). It may be that this form of SVMP PIII is peculiar to *B. arietans*, but it is possible that this form does exist in other SVMP PIII-rich vipers and because of the ease with which it can revert to monomer form during electrophoretic analysis, it may not have been characterised as a dimer. In many respects, this KEN SVMP PIII compares with the 68 and 75 kDa *B. arietans* SVMPs (BHRa and BHRb) characterised by (Omori-Satoh et al., 1995; Yamakawa et al., 1995). It is difficult to judge from their results what the native molecular weight might be, but these were able to degrade gelatins from a variety of collagens. It is likely that these BHRs (venom source unknown), our Kenyan gelatinase PIII and that observed in Currier et al., (2010), are all regional variants of the same protein.

Although these are preliminary biochemical studies, we can begin to relate the protease activities to haemotoxic features of puff adder envenoming. In particular, the potent, fast-acting laminin-degrading 21 kDa arilysin points to a possible role in the haemorrhagic action of the venoms which possess it. Some clinical studies have reported patients developing subcutaneous haemorrhage after cases of puff adder envenoming, but the region of origin of the snakes were not specified (Wakasugi et al., 2021, Husain et al., 2023). In the case of the gelatinolytic SVMP PIII, because of its potential ability to degrade fibrillar collagens at the dermal-epidermal junctions, this protease could contribute to blister formation at and around the bite site (Jiménez et al., 2008; Gutiérrez et al., 2018); this is a frequently observed feature of puff adder bites. At this stage, much of this is speculative and confirmation of this would require that these proteases are subject to further *in vitro* and *in vivo* studies; in the case of the possible haemorrhagic role of the 21 kDa arilysin. The exemplar for this would be the extensive characterisation that has been carried out on the haemorrhagic *Bothrops* SVMPs (Gutiérrez et al., 2016a, 2016b) from which we can extrapolate likely roles for the arilysins. Whatever its precise target(s), the work presented here shows that the potent 21 kDa arilysin surely plays a critical role in the tissue-damaging action of venoms from the puff adders which express it.

These SVMPs will undoubtedly underlie the mechanisms of coagulopathy, as is the case in other vipers. The pronounced fibrinogen-degrading activities of the 21 kDa arilysins will result in depletion of intact fibrinogen, suggesting that an anticoagulant effect may predominate, at least partly due to likely interference with the often-underestimated role of fibrinogen in platelet aggregation (Jackson, 2007). The glycosylated forms, particularly the two 30 kDa arilysins isolated from NGA and SDA venoms, were far less fibrinogenolytic. Compounding this issue is the marked variability in the levels of the glycosylated arilysins in the venoms that possess them. This is evident in Fig. S1, particularly in the three Namibian venoms, but we also observed a wide variety in its levels in the venom Nigerian (B) snakes; with one specimen having virtually no 30 kDa arilysin visible on SDS-PAGE (the two venoms in Fig. 1 lanes, 5 and 6 were the two venoms with the greatest amount). In contrast, in all those venoms that contained the 21 kDa arilysin, its level was markedly consistent, snake to snake and region to region. In an extensive study of the coagulotoxicity of venoms from bitis spp. Youngman et al., (2019), pronounced regional variations in the anticoagulant activity of *B. arietans* venoms was observed, with Tanzanian venoms being far more potent than those of West African snakes but also more so than that from nearby Kenya. Given the wide differences in the fibrinogen-degrading activity of the glycosylated and non-glycosylated forms of arilysin, it is tempting to speculate that the variations measured by Youngman et al. were due to differences in the types and levels of arilysin present in the venoms used.

The ability of the 21 kDa arilysins to completely degrade prothrombin and factor X also points to a complex effect of puff adder envenomation on the coagulation pathway. The inability to digest prothrombin or factor X in the specific manner required to activate them means that puff adder venom is highly unlikely to cause the consumptive coagulopathy as is seen in *Echis* spp., and, indeed, this is not an observed feature of puff adder envenoming. This general degradation of these key clotting factors will contribute to the overall anticoagulant effects of this venom, however.

## 5. Conclusion

The results presented here highlight the clinically significant regional variations in the SVMP activities of puff adder venoms. Variation was evident even within a single region, particularly the presence of the SVMP PIII gelatinase and the finding that NGA (B) venoms from snakes presently held at LSTM possessed the glycosylated 30 kDa arilysin, whereas stocks of NGA venom used in previous study contained the more potent non-glycosylated 21 kDa form. Considering this, it is perhaps not surprising that such a variety of clinical manifestations are observed following puff adder envenomation (Tianyi, 2024). It is reasonable to suggest that the severity of the reaction to envenomation will be related to whether the venom contains either the 21 kDa or the 30 kDa arilysin, with the 26 kDa Kenyan form being intermediate in this respect. This diversity in protease activity means that the well-documented issues with developing targeted therapeutics will be a particular problem for puff adder bites and, thus, this study illustrates the need for puff adder venoms from multiple geographic origins to ensure that conventional antivenom is effective across Africa. The work presented here is a good starting point to address these issues – by identifying the key venom SVMPs, in providing methods for their isolation prior to their use as immunogens for generation of toxin-specific antibodies and as the basis for experimental testing of small molecule therapeutics.

## Acknowledgements

We thank Herpetologists Paul Rowley and Edouard Crittenden at LSTM, and Geoffrey Kephah at KSRIC, for the maintenance and husbandry of the snake collection and the provision of venom samples.

The authors wish to thank the following for providing financial support to this study: Wellcome Trust grant 223619/Z/21/Z (to M.C.W., C.M.M., R.A.H., N.R.C.), Wellcome Trust grant 221712/Z/20/Z (to L-O.A., R.A.H., N.R.C.), The UK Foreign Commonwealth & Development Office grant 300341-115 (to R.A.H., N.R.C.), Department of Defense grant ID07200010-301-35 (to A.S.).

---

**Supplemental data** [Wilkinson 2025]

**Fig. S1.**
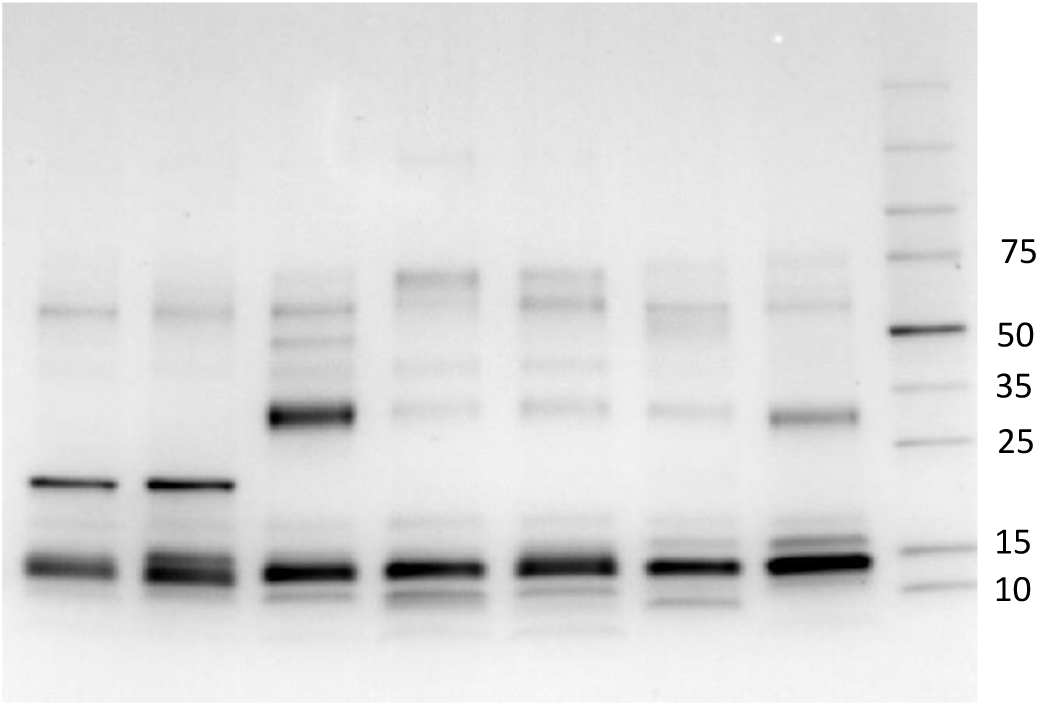
SDS-PAGE analysis of the venoms of individual *B. arietans* snakes from Nigeria, Namibia, South Africa and Saudi Arabia. The gel is 4-20% acrylamide (BioRad) and was run under reducing conditions. Eight μg of whole venom was loaded per lane and the gel and was stained using Coomassie Blue R250. The molecular weights of the markers (lane M, Thermo Broad Range) are indicated in kDa. The venoms shown, two samples of each, are 1, 2: Nigeria (NGA A); 3-5: Namibia (NAM); 6: South Africa (ZAF); 7: Saudi Arabia (SDA).

**Fig. S2.**
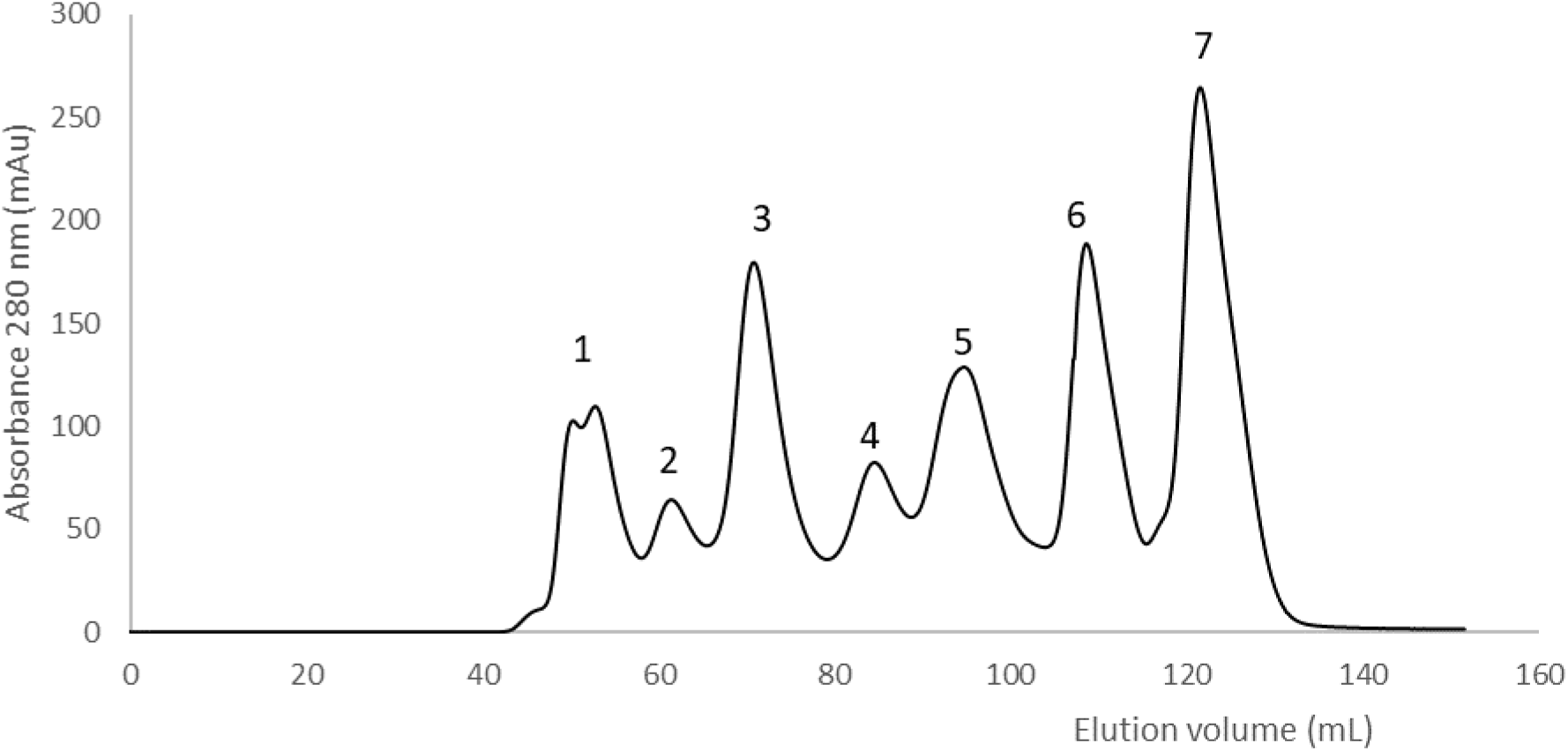
Size exclusion chromatography separation of TZA *B. arietans* venom. Whole venom (20 mg of TZA snake 006) was separated on a 120 mL column of Superdex 200HR equilibrated in 50 mM sodium phosphate pH 5.2. The flow-rate was 1.0 mL/min and elution was monitored at 280 nm.

**Fig. S3.**
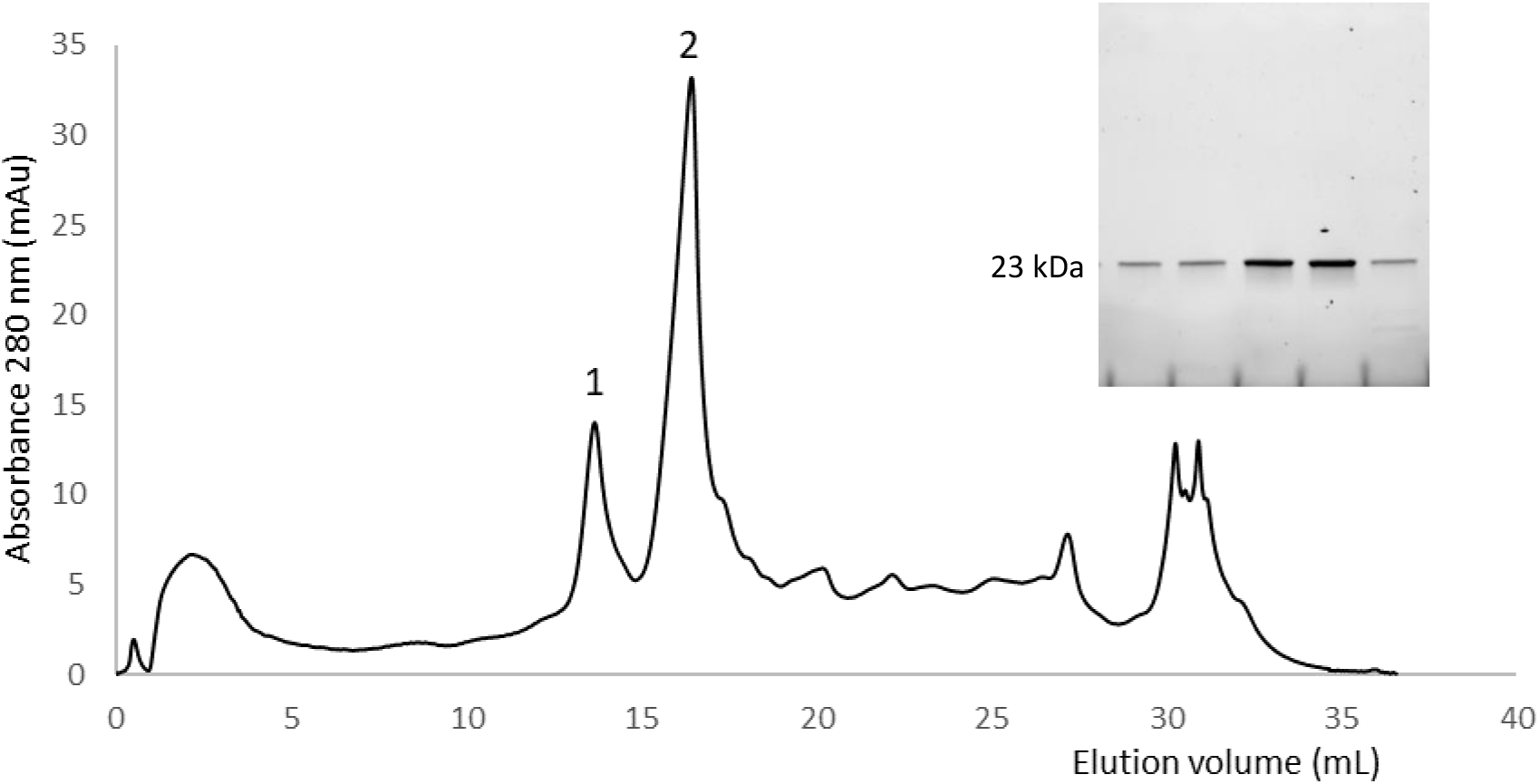
Cation exchange chromatography purification of TZA SVMPI. The proteins in peak 5 from SEC (Fig. S2) were applied to a 1 mL HiRes Capto S column equilibrated in 50 mM sodium phosphate pH 5.2. Elution was carried using a 25-column volume gradient of 0 - 0.25 M NaCl in 50 mM sodium phosphate pH 5.2. The flow rate was 0.6 mL/min and elution was monitored at 280 nm. Inset: SDS-PAGE analysis of the main fractions across peaks 1 and 2. The gel was visualised using the BioRad stain-free system.

**Fig. S4.**
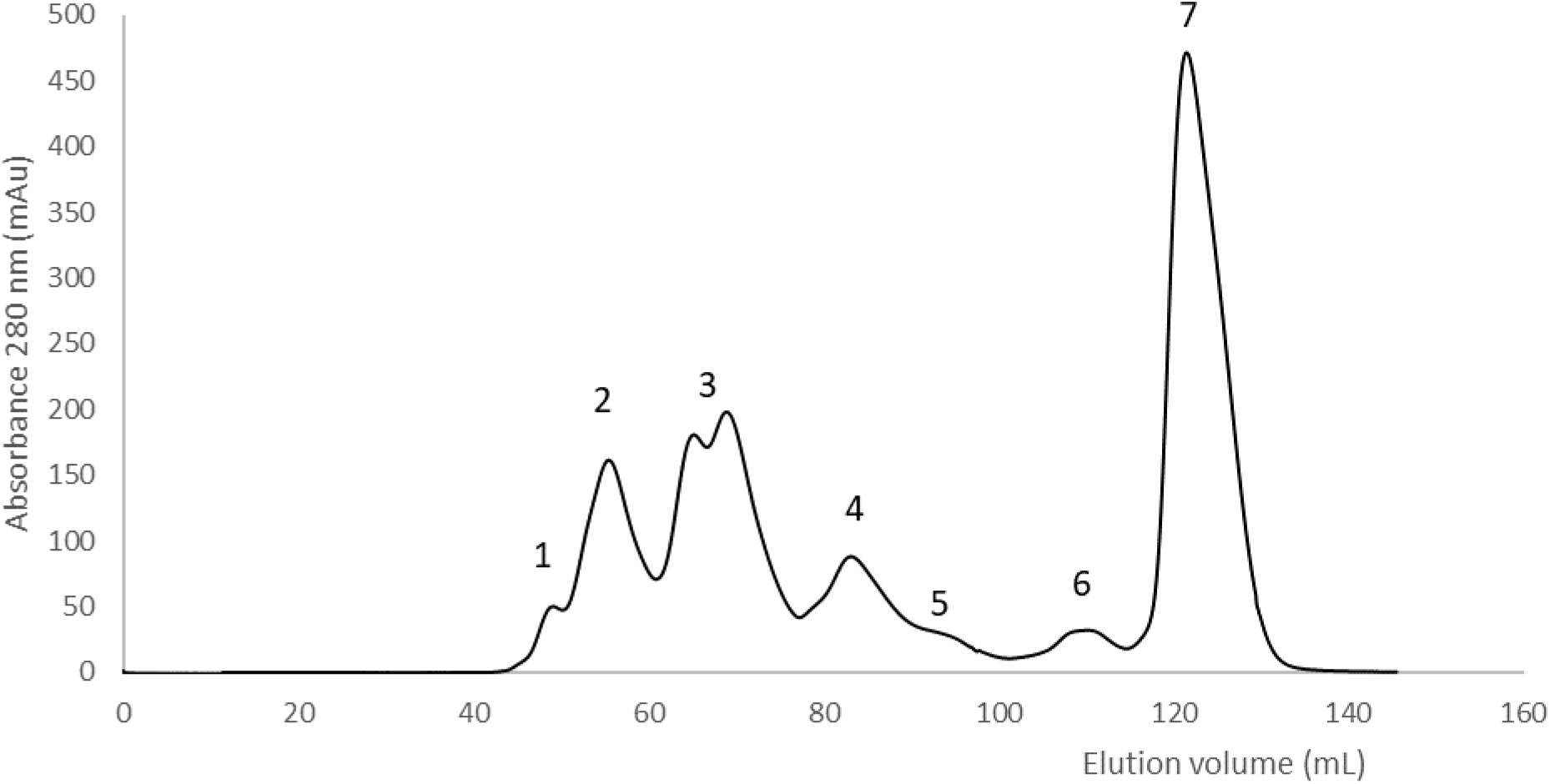
Size exclusion chromatography separation of NGA (B) *B. arietans* venom. Whole venom (20 mg NGA snake 011) was separated on a 120 mL column of Superdex 200HR equilibrated in 50 mM sodium phosphate pH 5.2. The flow-rate was 1.0 mL/min and elution was monitored at 280 nm.

**Fig. S5.**
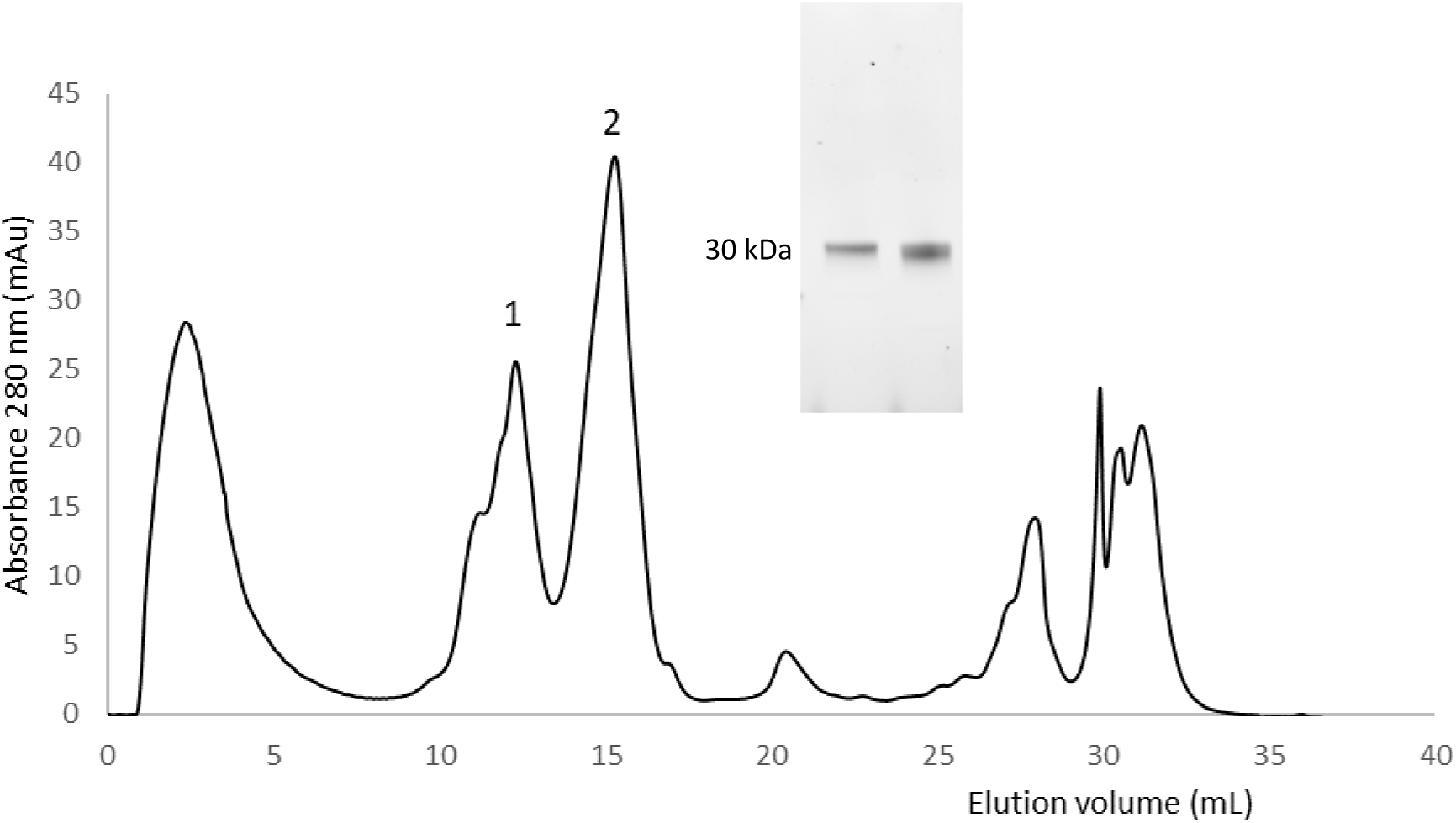
Cation exchange chromatography purification of NGA (B) SVMPI. The proteins in the first half peak 3 from SEC (Fig. 3S) were applied to a 1 mL HiRes Capto S column equilibrated in 50 mM sodium phosphate pH 5.2. Elution was carried using a 25-column volume gradient of 0 - 0.25 M NaCl in 50 mM sodium phosphate pH 5.2. The flow rate was 0.6 mL/min and elution was monitored at 280 nm. Inset: SDS-PAGE analysis of the main fractions in peaks 1 and 2. The gel was visualised using the BioRad stain-free system.

**Fig. S6.**
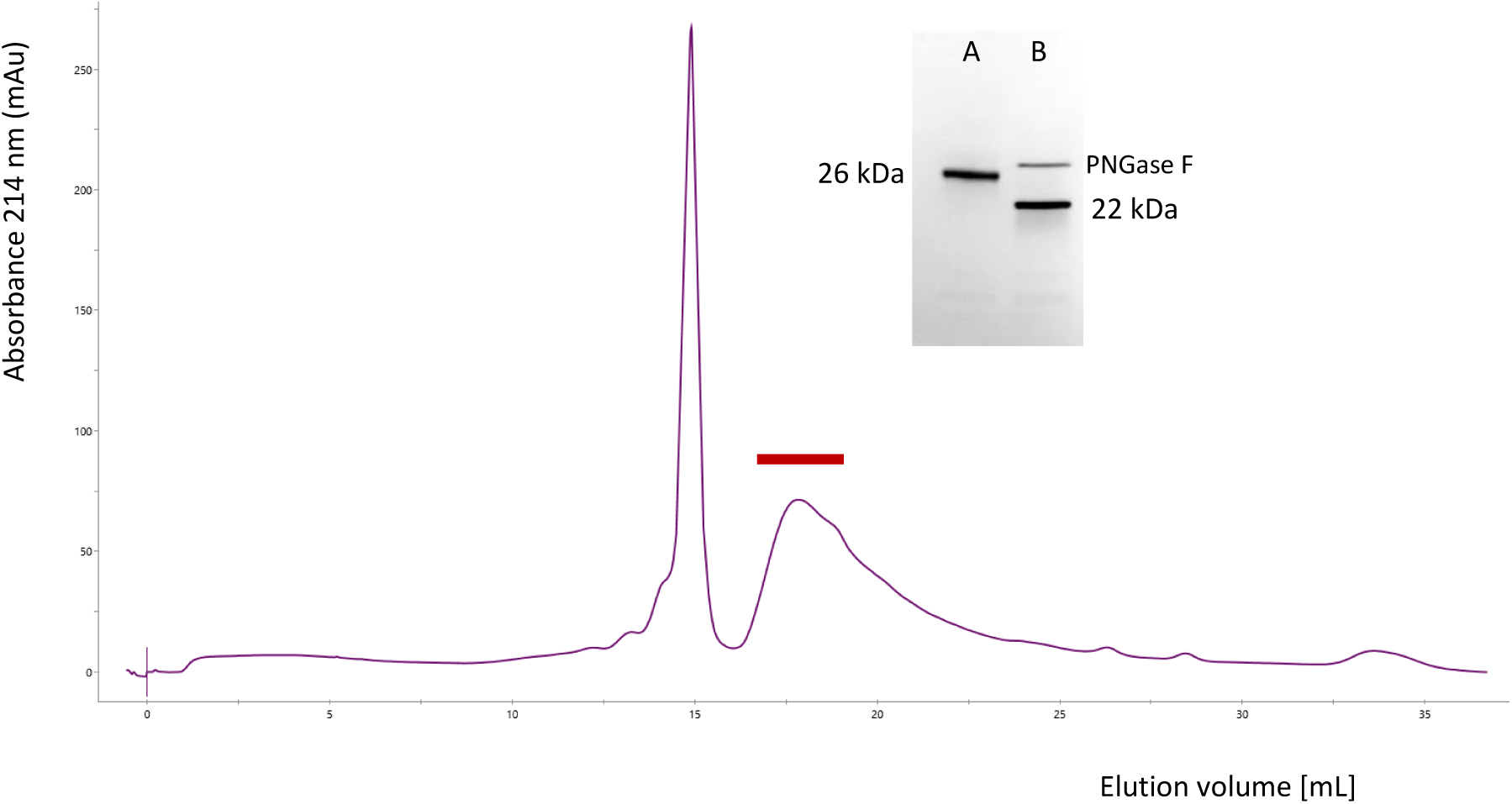
Cation exchange chromatography purification of Kenyan [KEN] SVMPI. The fractions from the initial cation exchange separation of whole venom (4.7 mL Capto SP column) containing the 32 kDa SVMPI proteins were desalted and applied to a 1.0 mL HiRes Capto S column equilibrated in 50 mM sodium phosphate pH 5.2. Elution was carried using a 25-column volume gradient of 0 - 0.50 M NaCl in 50 mM sodium phosphate pH 5.2. The flow rate was 1.0 mL/min and elution was monitored at 214 nm. Inset: SDS-PAGE analysis of the purified protein: A, main fractions [concentrated] from the SVMP PI peak (indicated with bar); B same after deglycosylation with PNGase F. The gel was stained with Coomassie Blue R250

**Fig. S7.**
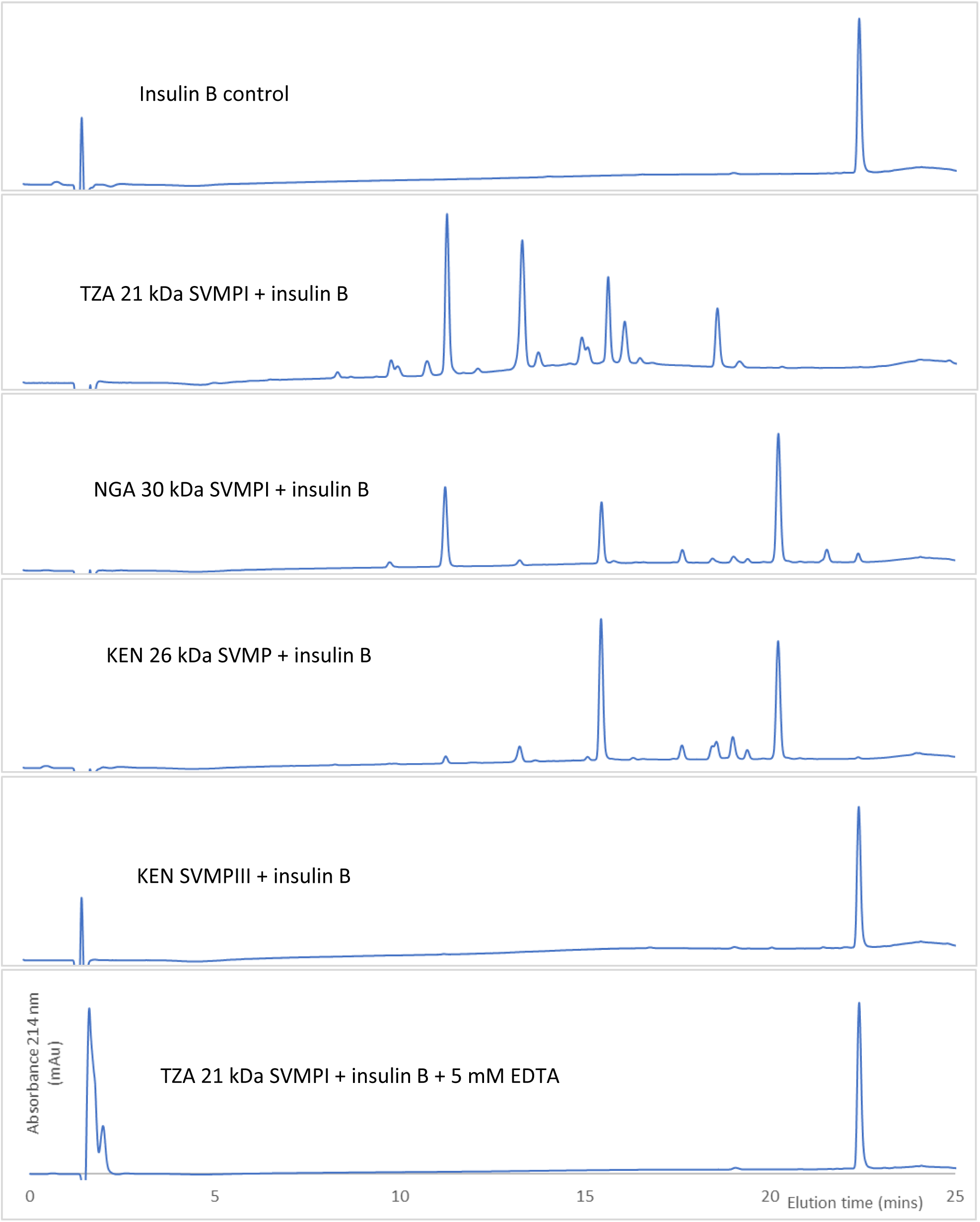
Insulin B degradation assay: Purified SVMPs. SVMPs were incubated at 37°C for 90 mins with oxidised insulin B chain at a ratio of 30:1 (w/w) insulin B: SVMP. An aliquot containing the equivalent of 1 ug insulin B was analysed by RP-HPLC with monitoring at 214 nm. The conditions for each are indicated in the inset. The result of an EDTA treated SVMP is shown for just one of the SVMPs (TZA SVMPI): all others were identical.

**Fig. S8.**
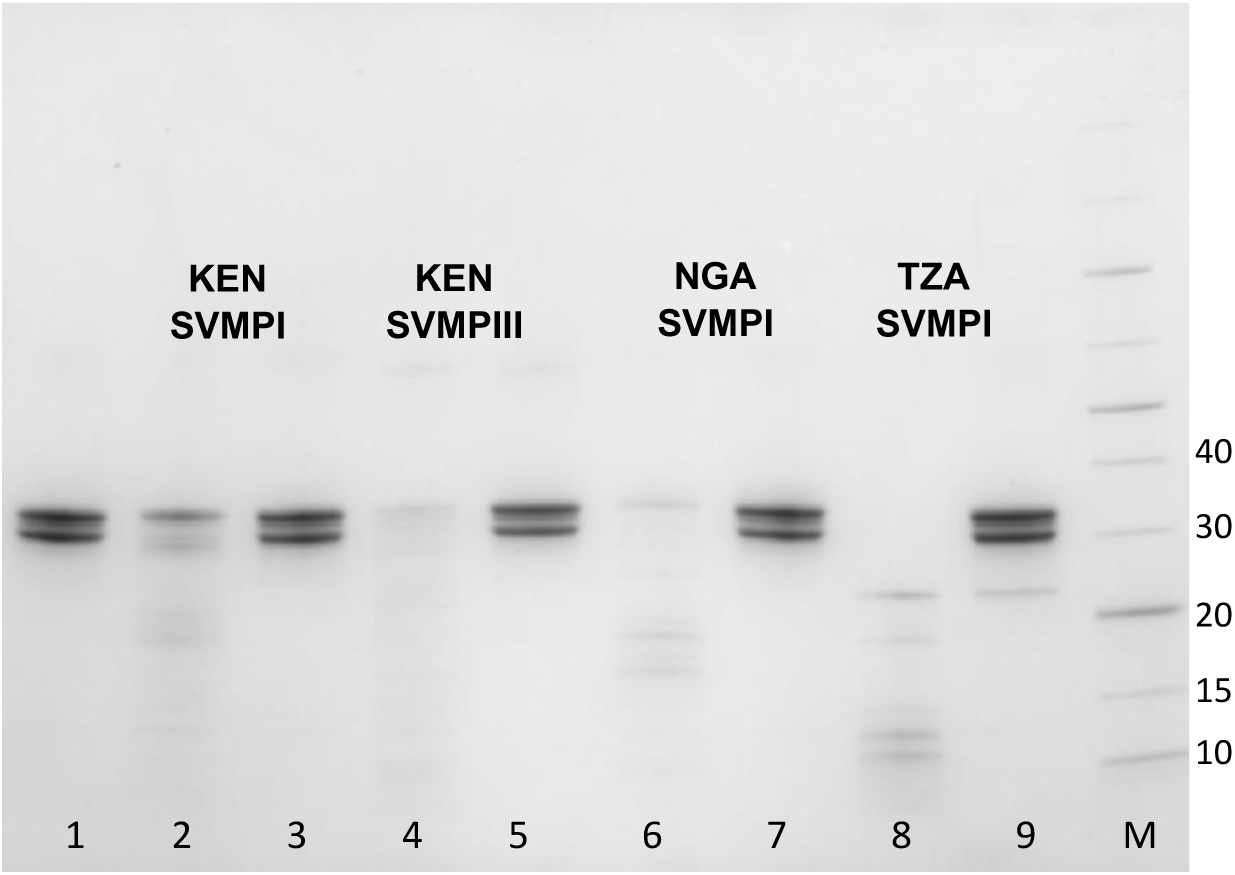
Casein degradation assay: Purified SVMPs. The purified SVMPs were incubated with bovine casein, either with or without 5 mM EDTA, for 90 minutes at 37 C. An aliquot of the reaction mix equivalent to 2 ug of casein was run on SDS-PAGE. The gel used was 4-20% acrylamide (BioRad) and was stained with Coomassie Blue R250. Lane 1, casein control; lanes 2 (no EDTA) and 3 (EDTA) KEN SVMP; lanes 4 (no EDTA) and 5 (EDTA) KEN SVMPIII; lanes 6 (no EDTA) and 7 (EDTA) NGA SVMPI; lanes 8 (no EDTA) and 9 (EDTA) TZA SVMPI; lane M, molecular weight markers (Thermo Page Ruler), indicated in kDa.

**Fig. S9.**
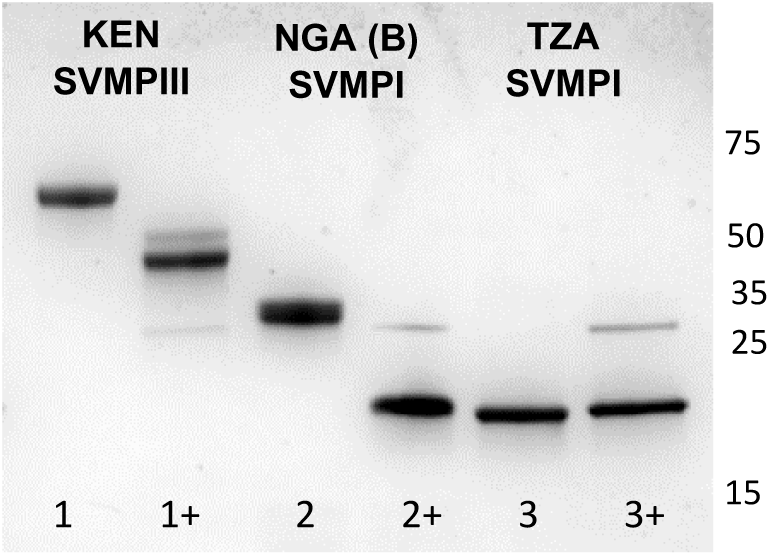
De-glycosylation of the main SVMPs used in this study. Samples were denatured then treated with PNGase F for 3 hours at 42°C. Lanes 1, 1+, KEN SVMPIII; lanes 2, 2+, NGA (B) SVMPI; lanes 3, 3+, TZA SVMPI. Lanes indicated + are the PNGase F-treated proteins. The sharp band at around 30 kDa in these lanes is PNGase F. The molecular weights of the markers (Promega Broad Range) are indicated in kDa on the right.

**Fig. S10.**
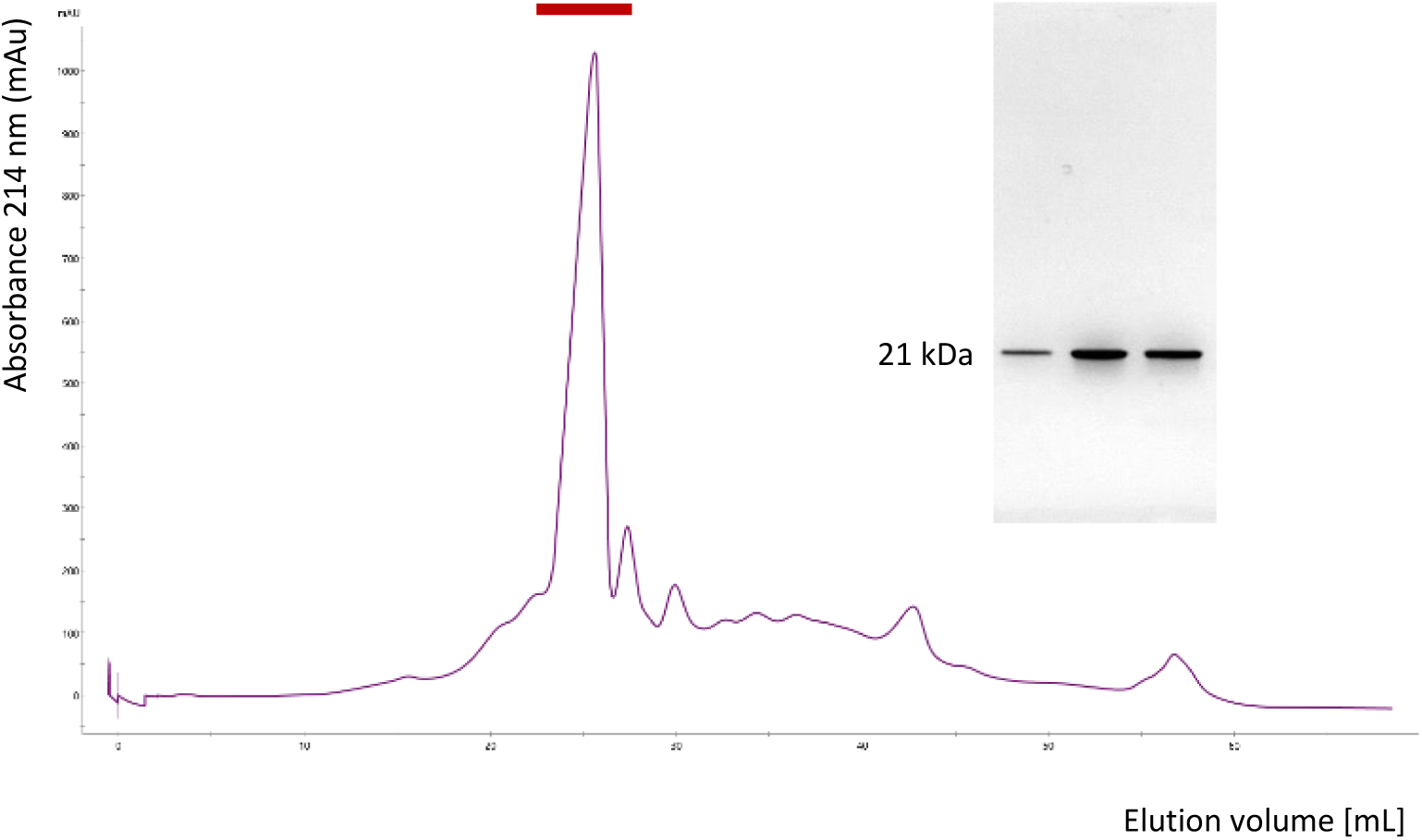
Cation exchange chromatography purification of GHA SVMPI. The fractions from SEC containing the 21 kDa SVMPI proteins were applied to a 4.7 mL Capto SP column equilibrated in 50 mM sodium phosphate pH 5.2. Elution was carried using a 10-column volume gradient of 0 - 0.40 M NaCl in 50 mM sodium phosphate pH 5.2. The flow rate was 0.6 mL/min and elution was monitored at 214 nm. Inset: SDS-PAGE analysis of the main fractions in the SVMP peak (indicated with bar). The gel was stained with Coomassie Blue R250.

**Fig. S11.**
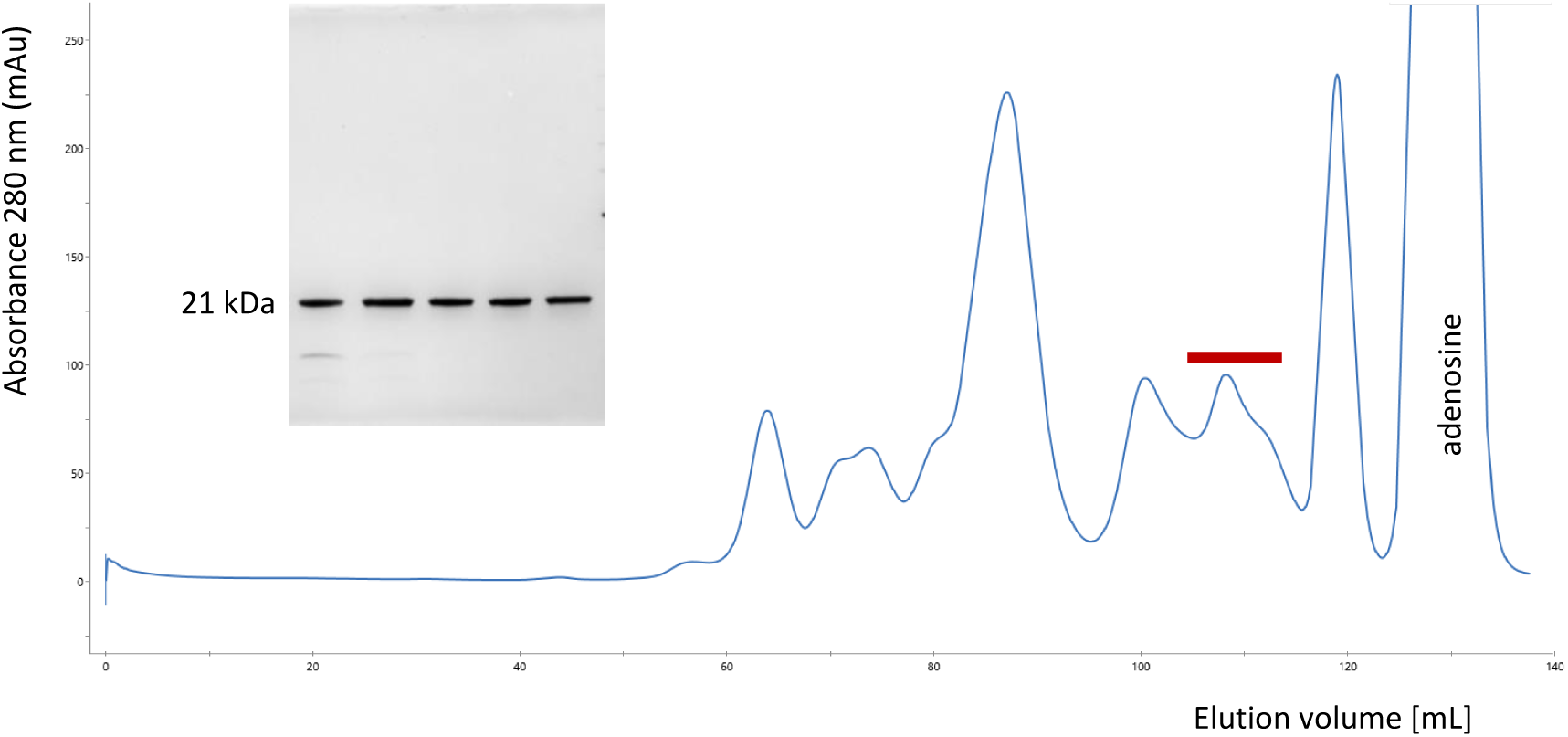
Size exclusion chromatography separation of Eswatini [ESW] *B. arietans* venom. Whole venom (25 mg in 1.2 mL) was separated on a 120 mL column of Superdex 200HR equilibrated in 50 mM sodium phosphate pH 5.2. The flow-rate was 1.0 mL/min and elution was monitored at 280 nm. Red bar indicates the peak containing the 21 kDa SVMP and the inset shows SDS-PAGE analysis of fractions within this peak. The gel was stained with Coomassie Blue R250.

**Fig. S12.**
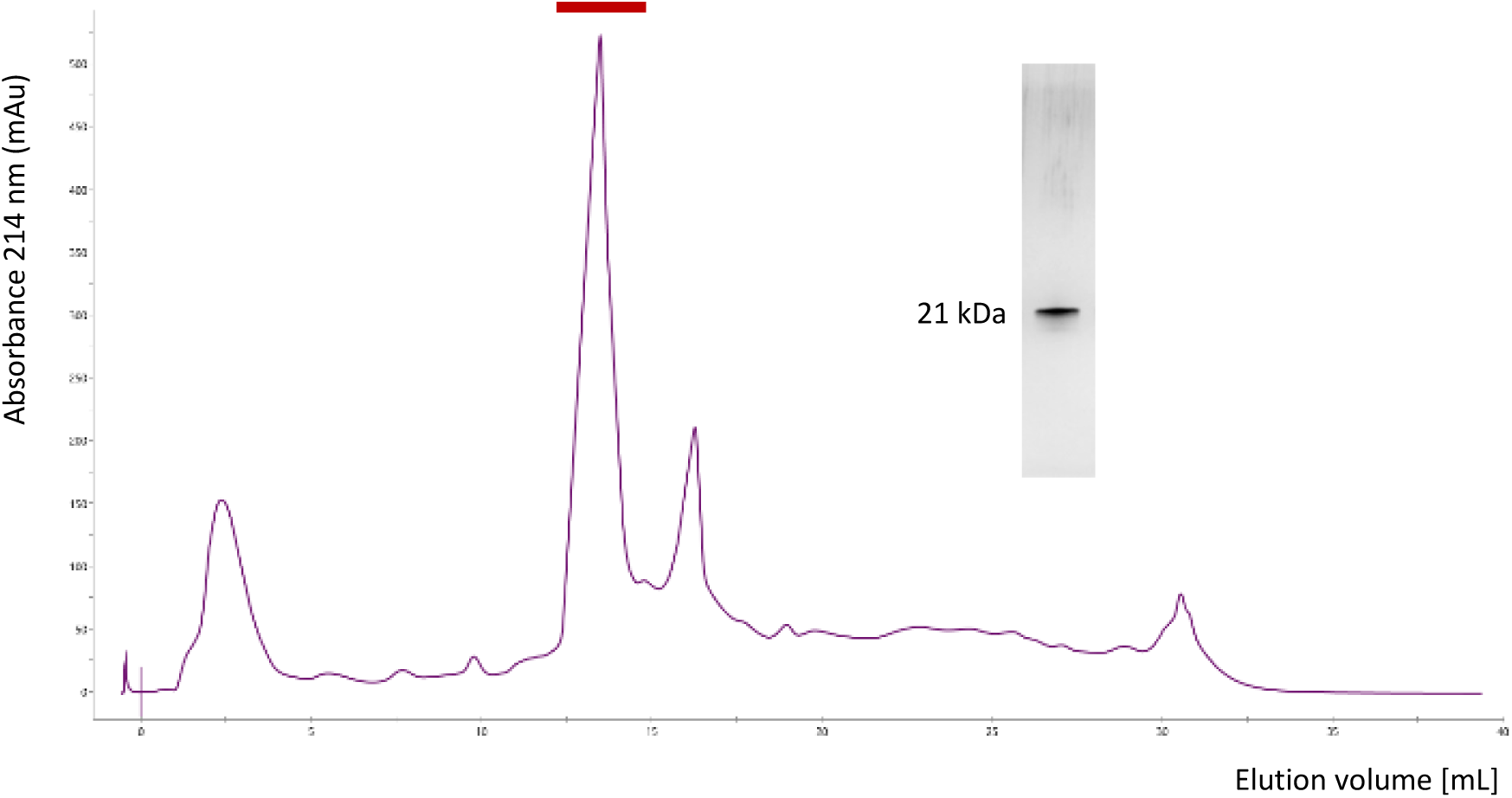
Cation exchange chromatography purification of NGA (A) SVMPI. The fractions from SEC containing the 21 kDa SVMPI proteins were applied to a 1.0 mL HiRes Capto S column equilibrated in 50 mM sodium phosphate pH 5.2. Elution was carried using a 25-column volume gradient of 0 - 0.50 M NaCl in 50 mM sodium phosphate pH 5.2. The flow rate was 1.0 mL/min and elution was monitored at 214 nm. Inset: SDS-PAGE analysis of the main peak, indicated with bar. The gel was stained with silver nitrate.

**Fig. S13.**
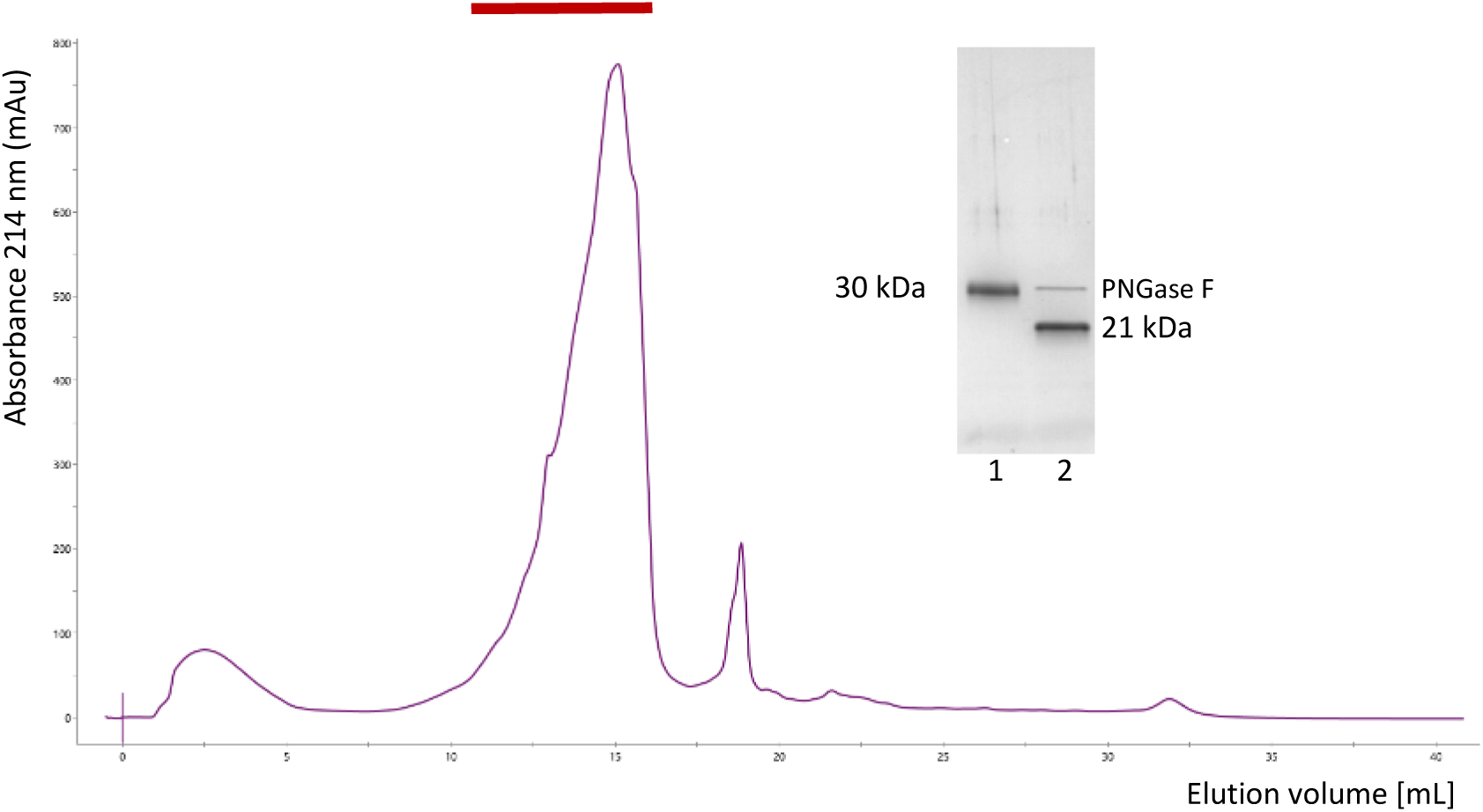
Cation exchange chromatography purification of Saudi Arabian (SDA) SVMP. The fractions from SEC containing the 30 kDa SVMPI proteins were applied to a 1.0 mL HiRes Capto S column equilibrated in 50 mM sodium phosphate pH 5.2. Elution was carried using a 25-column volume gradient of 0 - 0.60 M NaCl in 50 mM sodium phosphate pH 5.2. The flow rate was 0.5 mL/min and elution was monitored at 214 nm. Inset: SDS-PAGE analysis. Lane 1, protein from the main peak (indicated with bar); lane 2 same after PNGase F treatment. The gel was stained with Coomassie Blue R250.

**Fig. S14.**
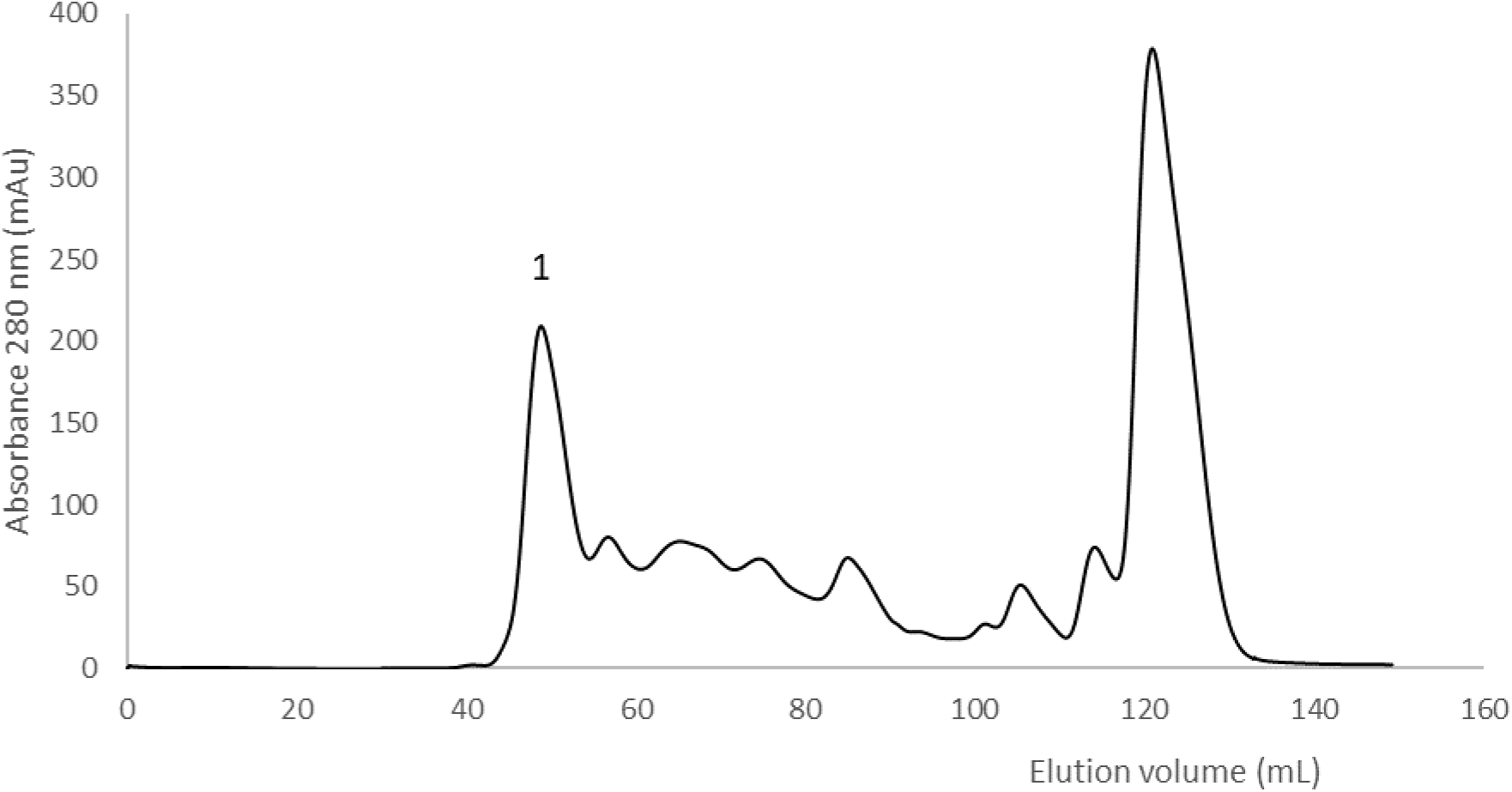
Size exclusion chromatography separation of KEN *B. arietans* venom. Whole venom (20 mg KEN015) was separated on a 120 mL column of Superdex 200HR equilibrated in 50 mM Tris-Cl pH 8.5. The flow-rate was 1.0 mL/min and elution was monitored at 280 nm.

**Fig. S15.**
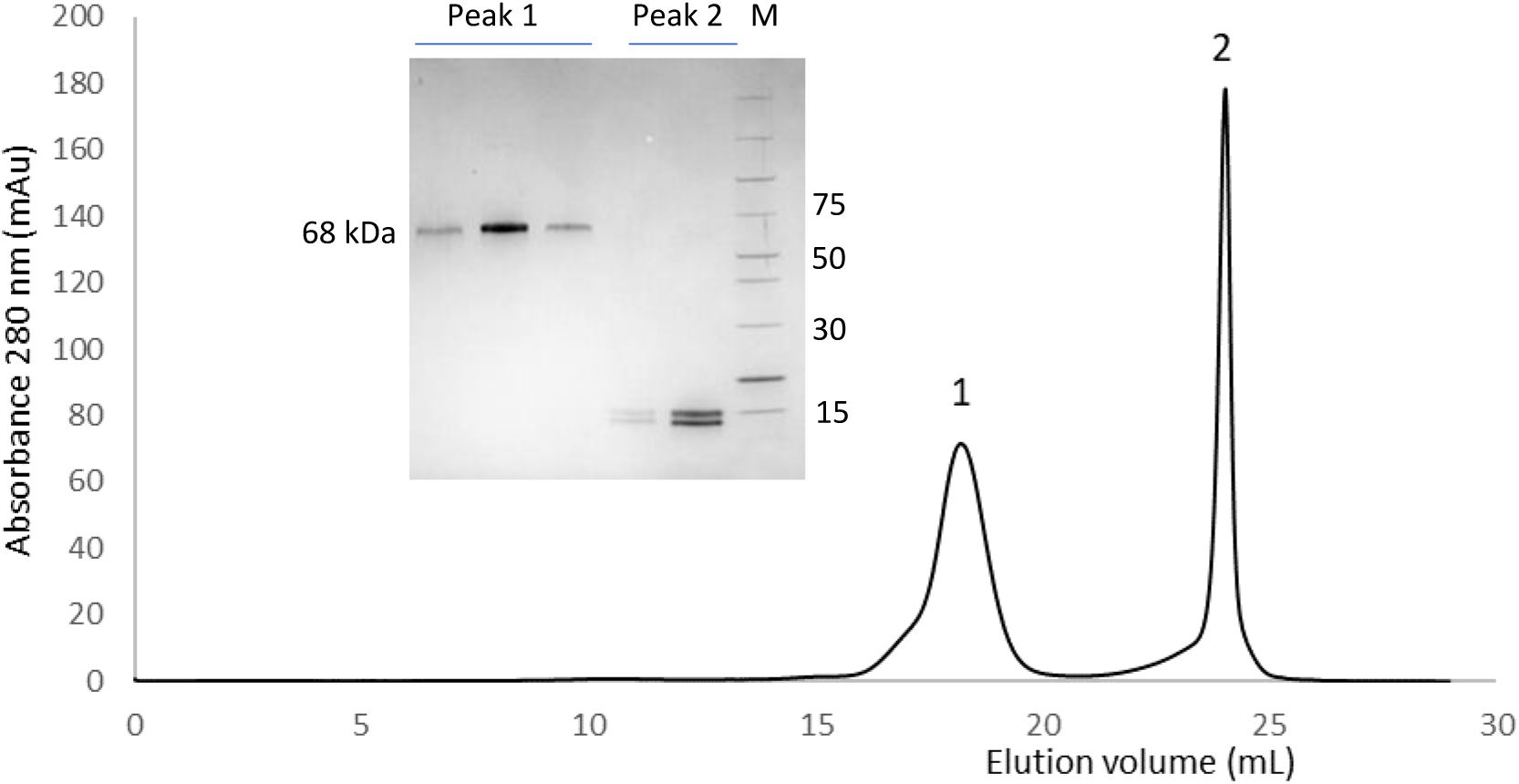
Anion exchange chromatography purification of KEN SVMPIII. The proteins containing the 60 kDa protein from SEC peak 1 (Fig. 8S) were applied directly to a 1 mL Mono Q column equilibrated in 50 mM Tris-Cl, pH 8.5. Elution was carried out using a 20-column volume gradient of 0-0.3 M NaCl in 50 mM Tris-Cl, pH 8.5. The column was operated at 0.6 mL/min and elution was monitored at 280 nm. Inset: SDS- PAGE analysis of fractions from the main peaks, lane M, molecular weight markers (Thermo Page Ruler), indicated in kDa. The gel was stained with Coomassie Blue R250

## References

Albulescu, L.-O., Hale, M.S., Ainsworth, S., Alsolaiss, J., Crittenden, E., Calvete, J.J., Evans, C., Wilkinson, M.C., Harrison, R.A., Kool, J., Casewell, N.R., 2020. Preclinical validation of a repurposed metal chelator as an early-intervention therapeutic for hemotoxic snakebite. Sci Transl. Med. 12 (542). 10.1126/scitranslmed.aay8314

Arlinghaus, F.T., Eble, J.A., 2012. C-type lectin-like proteins from snake venoms. Toxicon 60, 512–519. 10.1016/j.toxicon.2012.03.001

Baramova, E.N., Shannon, J.D., Bjarnason, J.B., Fox, J.W., 1989. Degradation of Extracellular Matrix Proteins by Hemorrhagic Metalloproteinases. Arch. Biochem. Biophys. 275, 63–71. 10.1016/0003-9861(89)90350-0

Casewell, N.R., Wagstaff, S.C., Wüster, W., Cook, D.A.N., Bolton, F.M.S., King, S.I., Pla, D., Sanz, L., Calvete, J.J., Harrison, R.A., 2014. Medically important differences in snake venom composition are dictated by distinct postgenomic mechanisms. PNAS 111, 9205–9210. 10.5061/dryad.1j292

Chevallet, M., Luche, S. and Rabilloud, T. 2006. Silver staining of proteins in polyacrylamide gels. Nature Protocols 1, 1852–1858. 10.1038/nprot.2006.288

Chippaux, J.P., 2011. Estimate of the burden of snakebites in sub-Saharan Africa: A meta-analytic approach. Toxicon 57, 586–599. 10.1016/j.toxicon.2010.12.022

Chung, M. C. M., Ponnudurai, G., Kataoka, M., Shimizu, S. and Tan N-H. 1996. Structural Studies of a Major Hemorrhagin (Rhodostoxin) from the Venom of Calloselasma rhodostoma. Arch. Biochem. Biophys. 325, 199–208. 10.1006/abbi.1996.0025

Currier, R.B., Harrison, R.A., Rowley, P.D., Laing, G.D., Wagstaff, S.C., 2010. Intra-specific variation in venom of the African Puff Adder (Bitis arietans): Differential expression and activity of snake venom metalloproteinases (SVMPs). Toxicon 55, 864–873. 10.1016/j.toxicon.2009.12.009

Dawson, C.A., Bartlett, K.E., Wilkinson, M.C., Ainsworth, S., Albulescu, L-O., Kazandijan, T Hall, S.R., Westhorpe, A., Clare, R., Wagstaff, S., Modahl, C.M., Harrison, R.A., Casewell, N.R. Intraspecific venom variation in the medically important puff adder (Bitis arietans): comparative venom gland transcriptomics, in vitro venom activity and immunological recognition by antivenom. bioRxiv 2024.03.13.584772; doi: 10.1101/2024.03.13.584772

Escalante, T., Shannon, J., Moura-da-Silva, A.M., María Gutiérrez, J., Fox, J.W., 2006. Novel insights into capillary vessel basement membrane damage by snake venom hemorrhagic metalloproteinases: A biochemical and immunohistochemical study. Arch. Biochem. Biophys. 455, 144–153. 10.1016/j.abb.2006.09.018

Fasoli, E., Sanz, L., Wagstaff, S., Harrison, R.A., Righetti, P.G., Calvete, J.J., 2010. Exploring the venom proteome of the African puff adder, Bitis arietans, using a combinatorial peptide ligand library approach at different pHs. J. Proteomics 73, 932–942. 10.1016/j.jprot.2009.12.006

Fling, S.P., Gregerson, S.F., 1986. Peptide and protein molecular weight determination by electrophoresis using a high-molarity Tris buffer system without urea. Anal. Biochem. 155, 83–88. 10.1016/0003-2697(86)90228-9

Fox, J.W., Serrano, S.M.T., 2008. Insights into and speculations about snake venom metalloproteinase (SVMP) synthesis, folding and disulfide bond formation and their contribution to venom complexity. FEBS Journal 275, 3016–3030 10.1111/j.1742-4658.2008.06466.x

Graham, R.L.J., McClean, S., O’Kane, E.J., Theakston, D., Shaw, C., 2005. Adenosine in the venoms from viperinae snakes of the genus Bitis: Identification and quantitation using LC/MS and CE/MS. Biochem. Biophys. Res. Commun. 333, 88–94. 10.1016/j.bbrc.2005.05.077

Gutiérrez, J.M., Escalante, T., Rucavado, A., Herrera, C., 2016a. Hemorrhage caused by snake venom metalloproteinases: A journey of discovery and understanding. Toxins 8, 93. 10.3390/toxins8040093

Gutiérrez, J.M., Escalante, T., Rucavado, A., Herrera, C., Fox, J.W., 2016b. A comprehensive view of the structural and functional alterations of extracellular matrix by snake venom metalloproteinases (SVMPs): Novel perspectives on the pathophysiology of envenoming. Toxins 8, 304. 10.3390/toxins8100304

Gutiérrez, J.M., Rucavado, A., Escalante, T., Herrera, C., Fernández, J., Lomonte, B., Fox, J.W., 2018. Unresolved issues in the understanding of the pathogenesis of local tissue damage induced by snake venoms. Toxicon 148, 123–131. 10.1016/j.toxicon.2018.04.016

Hite, L.A., Shannon, J.D., Bjarnason, J.B., Fox, J.W., 1992. Sequence of a cDNA clone encoding the zinc metalloproteinase hemorrhagic toxin e from Crotalus atrox: evidence for signal, zymogen and disintegrin-like structures. Biochemistry 31, 6203–6211. 10.1021/bi00142a005

Husain, Z., Wicaksono, A.C., Renault, A., Md Zhahir, S.S., Ismail, A.K., 2023. A case of fatal envenomation by a captive puff adder (Bitis arietans) in Malaysia. Toxicon 224. 10.1016/j.toxicon.2023.107023

Jackson, S.P., 2007. The growing complexity of platelet aggregation. Blood 109, 5087–5095. 10.1182/blood-2006-12-027698.

Jiménez, N., Escalante, T., Gutiérrez, J.M., Rucavado, A., 2008. Skin pathology induced by snake venom metalloproteinase: Acute damage, revascularization, and re-epithelization in a mouse ear model. Journal of Investigative Dermatology 128, 2421–2428. 10.1038/jid.2008.118

Juarez, P., Wagstaff, S. C., Oliver, J., Sanz, L., Harrison, R.A. and Calvete J. J. 2006. Molecular Cloning of Disintegrin-like Transcript BA-5A from a Bitis arietans Venom Gland cDNA Library:A Putative Intermediate in the Evolution of the Long-Chain Disintegrin Bitistatin. J. Mol. Evol. 63, 142–152. 10.1007/s00239-005-0268-z

Laemlli, U.K., 1970. Cleavage of Structural Proteins during the Assembly of the Head of Bacteriophage T4. Nature 227, 680–685. 10.1038/227680a0

Larkin, M.A., Blackshields, G., Brown, N.P., Chenna, R., McGettigan, P.A., McWilliam, H., Valentin, F., Wallace, I.M., Wilm, A., Lopez, R., Thompson, J.D., Gibson, T.J., Higgins, D.G., 2007. Clustal W and Clustal X version 2.0. Bioinformatics 23, 2947–2948. 10.1093/bioinformatics/btm404

Khin, T.Y., Pitts, M., Tongyoo, P., Rojnuckarin, P. and Wilkinson, M.C. 2017 Snake Venom Metalloproteinases and Their Peptide Inhibitors from Myanmar Russell’s Viper Venom. Toxins 9, 15. 10.3390/toxins9010015

Megale, A.A.A., Magnoli, F.C., Kuniyoshi, A.K., Iwai, L.K., Tambourgi, D. V., Portaro, F.C. V., da Silva, W.D., 2018. Kn-Ba: a novel serine protease isolated from Bitis arietans snake venom with fibrinogenolytic and kinin-releasing activities. Journal of Venomous Animals and Toxins including Tropical Diseases 24, 38. 10.1186/s40409-018-0176-5

Morita, T., 2005. Structures and functions of snake venom CLPs (C-type lectin-like proteins) with anticoagulant-, procoagulant-, and platelet-modulating activities. Toxicon 45, 1099–1114. 10.1016/j.toxicon.2005.02.021

Morita, T., Iwanaga, S., Suzuki, T., 1976. The mechanism of activation of bovine prothrombin by an activator isolated from Echis carinatus venom and characterization of the new active intermediates. J. Biochem. 79, 1089–1108. 10.1093/oxfordjournals.jbchem.a131150

Nikai, T., Momose, M., Okumura, Y., Ohara, A., Komori, Y., Sugihara, H., 1993. Kallidin-releasing enzyme from Bitis arietans (puff adder) venom. Arch. Biochem. Biophys. 307, 304–310. 10.1006/abbi.1993.1593

Olaoba, O.T., Karina dos Santos, P., Selistre-de-Araujo, H.S., Ferreira de Souza, D.H., 2020. Snake Venom Metalloproteinases (SVMPs): A structure-function update. Toxicon X 7, 100052. 10.1016/j.toxcx.2020.100052

Omori-Satoh Ap, T., Yamakawa, Y., Nagaoka, Y., Mebs, D., 1995. Hemorrhagic principles in the venom of Bitis arietans, a viperous snake. I. Purification and characterization. Biochim. et Biophys. Acta. 1246, 61–66. 10.1016/0167-4838(94)00170-L

Paixão-Cavalcante, D., Kuniyoshi, A.K., Portaro, F.C.V., da Silva, W.D., Tambourgi, D. V., 2015. African adders: Partial characterization of snake venoms from three Bitis species of medical importance and their neutralization by experimental equine antivenoms. PLoS Negl. Trop. Dis. 9. 10.1371/journal.pntd.0003419

Perkins, D.N., Pappin, D.J.C., Creasy, D.M., Cottrell, J.S., 1999. Probability-based protein identification by searching sequence databases using mass spectrometry data. Electrophoresis 20, 3551– 3567. http://doi.org/10.1002/(SICI)1522-2683(19991201)20:18<3551::AID-ELPS3551>3.0.CO;2-2

Sekoguchi, S., Nikai, T., Suzuki, Y., Sugihara, H., 1986. Kinin-releasing enzyme from the venom of Bitis arietans (puff adder), Biochim. et Biophvs. Acta 884, 502–509. 10.1016/0304-4165(86)90201-1

Serrano, S.M.T., 2013. The long road of research on snake venom serine proteinases. Toxicon 62, 19–26. 10.1016/j.toxicon.2012.09.003

Shimokawa, K., Jia, L.G., Wang, X.M. and Fox J.W. 1996. Expression, Activation, and Processing of the Recombinant Snake Venom Metalloproteinase, Pro-Atrolysin E. Arch. Biochem. Biophys. 335, 283–294. 10.1006/abbi.1996.0509

Strydom, D.J., Joubert, F.J., Howard, N.L.,1986. Chemical studies on Protease A of Bitis arietans (puff adder) Venom. Toxicon 24, 247–257. 10.1016/0041-0101(86)90150-9

Tan, N-H., Ponnudurai, G. and Chung, M. C. M. (1997) Proteolytic Specificity of Rhoddostoxin, the Major Hemorrhagin of Calloselasma Rhodostoma Venom. Toxicon. 35, 979–984. 10.1016/S0041-0101(96)00186-9

Tianyi, F-L., Ngari, C., M.C., Parkurito, S., Chebet, E., Mumo, E., Trelfa, A., Otundo, D., Crittenden, E., Kephah, G.M., Harrison, R.A., Stienstra, Y., Casewell, N.R., Lalloo, D.G., Oluoch, G.O. Clinical features of puff adder envenoming: case series of Bitis arietans snakebites in Kenya and a review of the literature. 2024. https://medrxiv.org/cgi/content/short/2024.05.31.24308288

Toth, M., Sohail, A., Fridman, R., 2012. Assessment of gelatinases (MMP-2 and MMP-9) by gelatin zymography. Methods in Molecular Biology 878, 121–135. 10.1007/978-1-61779-854-2_8

Wagstaff, S.C., Favreau, F., Cheneval, O., Laing, G.D., Wilkinson, M.C., Miller, R.L., Stocklin, R. and Harrison, R.A., 2008. Molecular characterisation of endogenous snake venom metalloproteinase inhibitors. Biochem. Biophys. Res. Commun. 365, 650–656. 10.1016/j.bbrc.2007.11.027

Van Der Walt, S.J., Joubert, F.J., 1971. Studies on puff adder (Bitis arietans) venom-I. Purification and properties of Protease A. Toxicon 9, 153–161. 10.1016/0041-0101(71)90009-2

Van Der Walt, S.J., Joubert, F.J., 1972. studies on puff adder (Bitis arietans) venom-II. Specificity of Protease A. Toxicon 10, 341–349. 10.1016/0041-0101(72)90056-6

Wakasugi, M., Kawagishi, T., Hatano, T., Shibuya, T., Kuwano, H., Matsui, K., 2021. Case report: Treatment of a severe Puff adder snakebite without antivenom administration. American Journal of Tropical Medicine and Hygiene 105, 525–527. 10.4269/ajtmh.21-0291

Warrell, D.A., Ormerod, L.D., Davidson, N.M., 1975. Bites by puff-adder (Bitis arietans) in Nigeria, and Value of antivenom. Br. Med. J. 4, 697–700. 10.1136/bmj.4.5998.697

Yamada, D., Sekiya, F., Morita, T., 1996. Isolation and Characterization of Carinactivase, a novel prothrombin activator in Echis carinatus venom with a unique catalytic mechanism. J. Biol. Chem. 271, 5200–5207. 10.1074/jbc.271.9.5200

Yamakawa, Y., Omori-Satoh, T., Mebs, D., 1995. Hemorrhagic principles in the venom of Bitis arietans, a viperous snake. II. Enzymatic properties with special reference to substrate specificity. Biochim. et Biophys. Acta 1247, 17–23. 10.1016/0167-4838(94)00171-C

